# Unanchored ubiquitin chains promote the non-canonical inflammasome via UBXN1

**DOI:** 10.1101/2024.10.30.621131

**Authors:** Duomeng Yang, Jason G. Cahoon, Tingting Geng, Chengliang Wang, Andrew G. Harrison, Evelyn Teran, Yanlin Wang, Anthony T. Vella, Vijay A. Rathinam, Jianbin Ruan, Penghua Wang

**Affiliations:** Department of Immunology, School of Medicine, UConn Health, Farmington, CT 06030, USA; Department of Medicine, School of Medicine, UConn Health, Farmington, CT 06030, USA; Harvard Medical School, Boston Children’s Hospital, Division of Immunology, Boston, MA, USA

**Keywords:** Inflammasome, UBXN1, Caspase-4/11, Ubiquitination, Unanchored ubiquitin chain

## Abstract

Ubiquitination is a major posttranslational covalent modification that regulates numerous cellular processes including inflammasome signaling. Cells also contain unanchored ubiquitin chains (polyUb) that bind protein targets non-covalently, but their physiological functions in immunity have been appreciated only recently. Here, we report that ubiquitin regulatory x domain-containing protein 1 (UBXN1) activates the noncanonical inflammasome via unanchored Lysin (K) 48- or 63-linked polyUb. UBXN1 deficiency impairs the activation of caspase-4/11, secretion of inflammasome-dependent cytokines and pyroptosis in response to intracellular lipopolysaccharide (LPS). UBXN1-deficient mice are protected from LPS- and cecal-ligation-and-puncture-induced sepsis, evidenced by reduced mortality and systemic inflammation, compared to UBXN1-sufficient littermates. Mechanistically, UBXN1 together with unanchored K48/63-linked polyUb bind caspase-4/11, the intracellular sensors of LPS, and promote their assembly and activation. Depleting cellular unanchored polyUb with recombinant ubiquitin-specific proteinase 5 (USP5) reduces UBXN1 binding to caspase-4/11 and inflammasome signaling, while USP5 inhibitors enhance pyroptosis in an UBXN1-dependent manner. Thus, this study identifies a critical UBXN1-dependent posttranslational mechanism involved in noncanonical inflammasome activation and UBXN1 as a potential therapeutic target for sepsis and advances a fundamental understanding of unanchored polyUb biology.

## INTRODUCTION

Inflammasomes are large cytosolic multiprotein complexes formed in response to infections and cellular stresses, leading to auto-activation of inflammatory caspases (human caspase-1, -4, -5, murine caspase-1, -11), secretion of inflammatory mediators [Interleukin (IL) -1 and -18)] and pyroptosis, a form of cell death (*1*). Canonical inflammasomes are assembled by one of the NLRs [Nucleotide-binding and oligomerization domain (NOD), leucine-rich repeat (LRR)- containing receptors (NLRs)], AIM2 (absent in melanoma 2) or Pyrin, upon binding by a microbial ligand or cellular danger signal. They then oligomerize, recruit the adaptor ASC (Apoptosis-associated speck-like protein containing a caspase recruitment domain) (optional), then pro-caspase-1, which undergoes self-cleavage into an active form. Active caspase-1 then proteolytically converts pro-IL-1β and pro-IL-18 into their biologically active, secreted forms (*2*). The non-canonical inflammasomes are activated by direct binding of a bacterial endotoxin, lipopolysaccharide (LPS), to human caspase-4/5 (equivalent to murine caspase-11), which preferably cleaves pro-IL-18. Regardless of their preference for different cytokines, all the inflammatory caspases cleave Gasdermin D (GSDMD) into an active N-terminal fragment that forms pores in the plasma membrane, leading to cell death (*3*).

Although inflammasomes are critical for the control of many microbial infections, hyperactive inflammasome signaling may lead to sepsis as well as autoinflammatory / autoimmune conditions (*4*). For example, the caspase-11 inflammasome is the main driver of LPS and polymicrobial sepsis in mice (*5–8*), and caspase-4 inflammasome-mediated pyroptosis facilitates immunocoagulation and associates with organ damage in human sepsis and septic shock (*9–11*). Therefore, inflammasome signaling is subjected to tight checks-and-balances on both the transcriptional and posttranslational levels. In particular, ubiquitination is one of the most important posttranslational regulations for numerous immune signaling pathways, including the canonical NLRP3 inflammasome (*12*). Many ubiquitin E3 ligases have been identified to either inhibit NLRP3 signaling via degradation (*13–17*) or promote NLRP3-caspase-1 activation via K63-linked polyubiquitination (*18–20*). However, little is known about the function of ubiquitination in the non-canonical inflammasomes (*21*). Only one E3 ligase, neural precursor cell expressed developmentally down-regulated protein 4 (NEDD4), was reported to induce K48-linked polyubiquitination and subsequent degradation of caspase-11 (*22*). Whether ubiquitination is required for the activation of caspase-4 and 11 inflammasomes remains unknown.

By immunoprecipitation coupled with mass spectrometry, we identified ubiquitin regulatory x domain-containing protein 1 (UBXN1) as a caspase-4 interacting protein. UBXN1 deficiency leads to impaired activation of caspase-4/11 and GSDMD, secretion of inflammasome-dependent cytokines and pyroptosis in response to intracellular LPS in HeLa cells and primary mouse macrophages. Consequently, UBXN1-deficient mice are protected from LPS- and cecal-ligation-and-puncture (CLP)-induced sepsis, evidenced by reduced mortality and systemic inflammation, compared to UBXN1-sufficient littermates. Mechanistically, UBXN1 promotes unanchored Lysine 48/63-linked polyubiquitin chains (K48/63-Ub) binding to caspase-4/11, thus facilitating their assembly and activation. Therefore, during sepsis targeting UBXN1 and unanchored polyUb may provide rapid relief from a cytokine storm by circumventing inflammasome activation.

## RESULTS

### UBXN1 is a potent binder of caspase-4/11

Ubiquitination is a major posttranslational regulation for immune signaling, however, little is known about its function in the non-canonical inflammasomes (*3*). To identify ubiquitination-related interactors of caspase-4 in an unbiased manner, we expressed FLAG-CASP4 in HeLa cells (a cancerous human epithelial cell line suitable for studying the non-canonical inflammasome), performed immunoprecipitation (IP) with an anti-FLAG antibody, and identified caspase-4-bound endogenous proteins by mass spectrometry. We identified 161 putative caspase-4 binders, however, only five are involved in ubiquitination, including two UBXN family members (UBXN1 and UBXN3B), two E3 ligases [STUB1 (STIP1 homology and U-box containing protein 1) and HUWE1 (HECT, UBA and WWE domain containing 1)], and ubiquitin (**Fig.1a, b**). To examine their role in caspase-4 activation, we generated stable polyclonal UBXN1, STUB1, HUWE1, and CASP4 knockout HeLa cells by CRISPR-Cas9 (Supplemental **Fig.S1a**). The UBXN3B phenotype was investigated with bone marrow-derived macrophages (BMDMs) derived from our previously generated tamoxifen-inducible knockout mouse model (*23–25*). We noted an obvious difference in cell death between *UBXN1*^-/-^ and its corresponding WT cells following LPS transfection, like *CASP4*^-/-^ (**Fig. 1c**; Supplemental **Fig.S1b**). Thus, we sought to explore UBXN1 function in the caspase-4 inflammasome. Next, we validated the caspase-4-UBXN1 interaction by co-immunoprecipitating endogenous caspase-4 and Myc-UBXN1 with an anti-MyC antibody (**Fig.1d**). To estimate the caspase-4-UBXN1 binding capacity, we overexpressed FLAG-APEX-CASP4 and Myc-UBXN1 together and immunoprecipitated FLAG-APEX-CASP4. We observed that nearly an equal amount of Myc-UBXN1 was co-immunoprecipitated (**Fig.1e**), indicating a strong interaction. We confirmed that caspase-11-the murine counterpart of human caspase-4-was also able to pull down Myc-UBXN1, but human caspase-1 could not (**Fig.1f**), indicating that UBXN1 specifically targets the non-canonical inflammasome. Next, to validate the above immunoprecipitation results, we investigated endogenous UBXN1-caspase-4/11 colocalization by confocal microscopy. In resting cells (mock), caspase-4/11 was largely diffuse throughout the cytoplasm, while UBXN1 was dispersed in the cytoplasm and nucleoplasm. Notably, following LPS transfection, UBXN1 was significantly colocalized with caspase-4/11 to large punctate structures, typical of inflammasomes, in HeLa and mouse primary macrophages respectively (**Fig.1g, h**). These data suggest that UBXN1 is a component of the caspase-4/11 non-canonical inflammasome complex.

**Fig.1.**
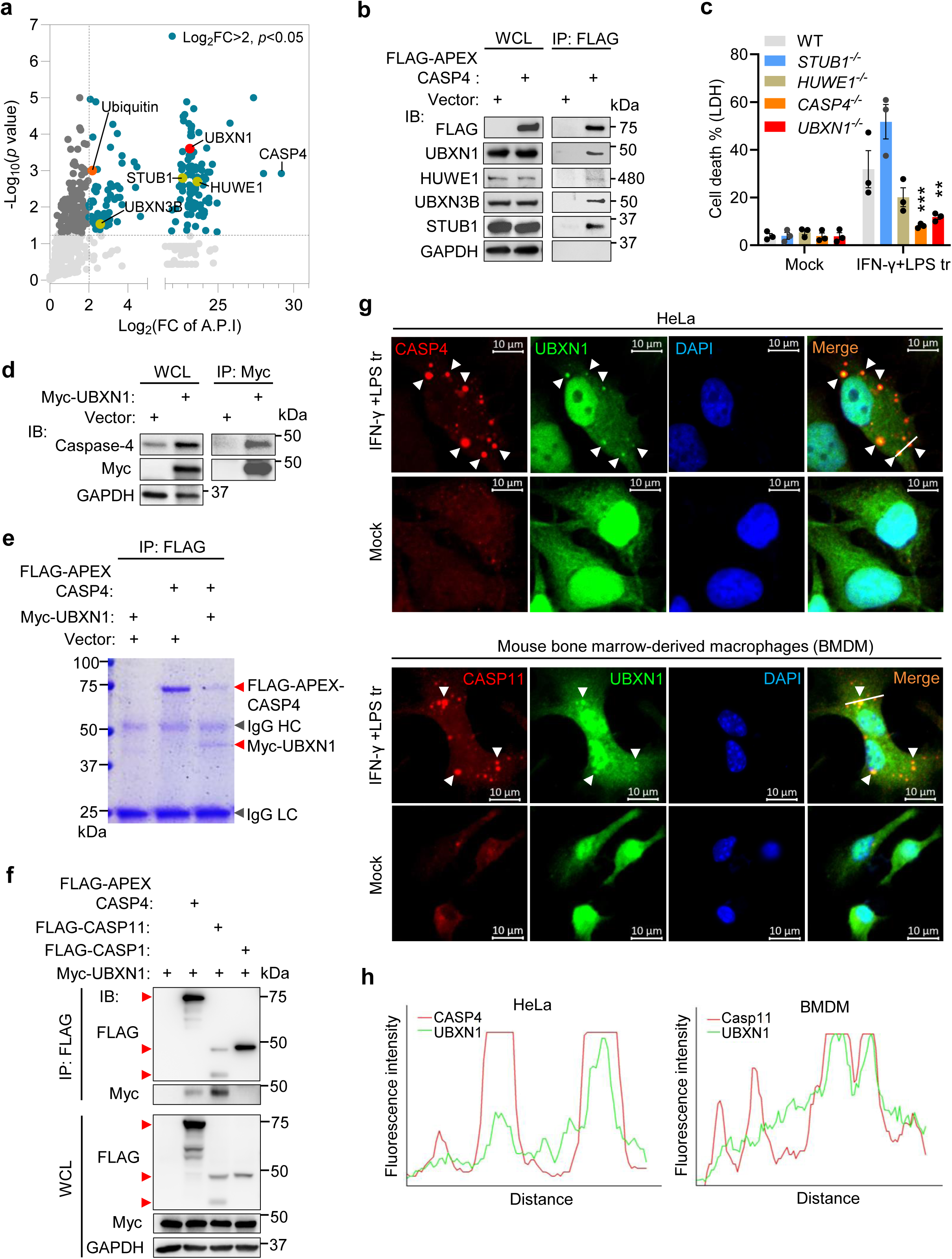
UBXN1 binds caspase-4/11. (**a**) A scatter plot showing the endogenous proteins co-immunoprecipitated (IP) with FLAG-CASP4 and identified by mass spectrometry. Proteins involved in ubiquitination are labeled. Dark green dots indicate the proteins with the Log_2_ fold-change (FC) of Average Precursor Intensity (A.P.I)□>□2, *p*<0.05 by two-tailed Student’s t-test with Benjamini–Hochberg. The Log_2_ FC is calculated as Log_2_ (A.P.I of a protein from the FLAG-CASP4 / Vector group). (**b**) Co-IP of FLAG-APEX-caspase-4 and endogenous interactors with an anti-FLAG antibody from HeLa cells transfected with a FLAG-APEX-CASP4 plasmid. Shown are the immunoblots (IB) of indicated proteins. WCL, whole cell lysate. (**c**) The percent cell death (LDH release) of HeLa cells primed with 10 ng/mL of human IFN-γ for 12 h followed by transfection with 1 μg/mL LPS (LPS tr) for 5 h. Mock are untreated controls. Data are presented in mean ± S.E.M, each symbol represents a biological replicate, \*\**p*<0.01, \*\*\**p*<0.001 by Two-Way ANOVA, Dunnet comparison test. (**d**) Co-IP of Myc-UBXN1 and endogenous caspase-4 with an anti-Myc antibody from HeLa cells transfected with a Myc-UBXN1 plasmid. (**e**) Co-IP of FLAG-APEX-caspase-4 and Myc-UBXN1 with an anti-FLAG antibody from HeLa cells transfected with a FLAG-APEX-CASP4 and Myc-UBXN1 plasmid. Shown is Coomassie blue staining of an SDS-PAGE gel. IgG HC, immunoglobin heavy chain; LC, light chain. (**f**) Co-IP of FLAG-caspases and Myc-UBXN1 with an anti-FLAG antibody from HEK293T cells transfected with a FLAG-CASP and Myc-UBXN1 plasmid. The red arrow heads indicate desired protein bands. The recombinant caspase-4 has FLAG and APEX tags. FLAG-caspase-11 shows in two bands. (**g**) Immunofluorescent confocal images in HeLa cells and mouse bone marrow-derived macrophages (BMDM) that were primed with IFN-γ and transfected with LPS. Mock, transfection reagent only. DNA was counterstained by DAPI. Arrows point to colocalizations. Scale bar: 10 µm. Lines indicate the plot profiles for (**h**). (**h**) Histogram analyses of the fluorescence intensity distribution in (**g**).

### UBXN1 is crucial for the non-canonical inflammasome signaling

Next, we in-depth characterized the role of UBXN1 in non-canonical inflammasome signaling. We generated multiple monoclonal knockout lines, and recapitulated the reduction of pyroptosis (Supplemental **Fig.S2**, **Fig.2a**). Consistent with the cell death assay, both active caspase-4 p32 and GSDMD-N fragments appeared in WT cells after LPS transfection, while their levels were greatly reduced in *UBXN1*^-/-^ cells (**Fig.2b**) as were maturation and secretion of IL-18 (**Fig.2c**).

**Fig. 2.**
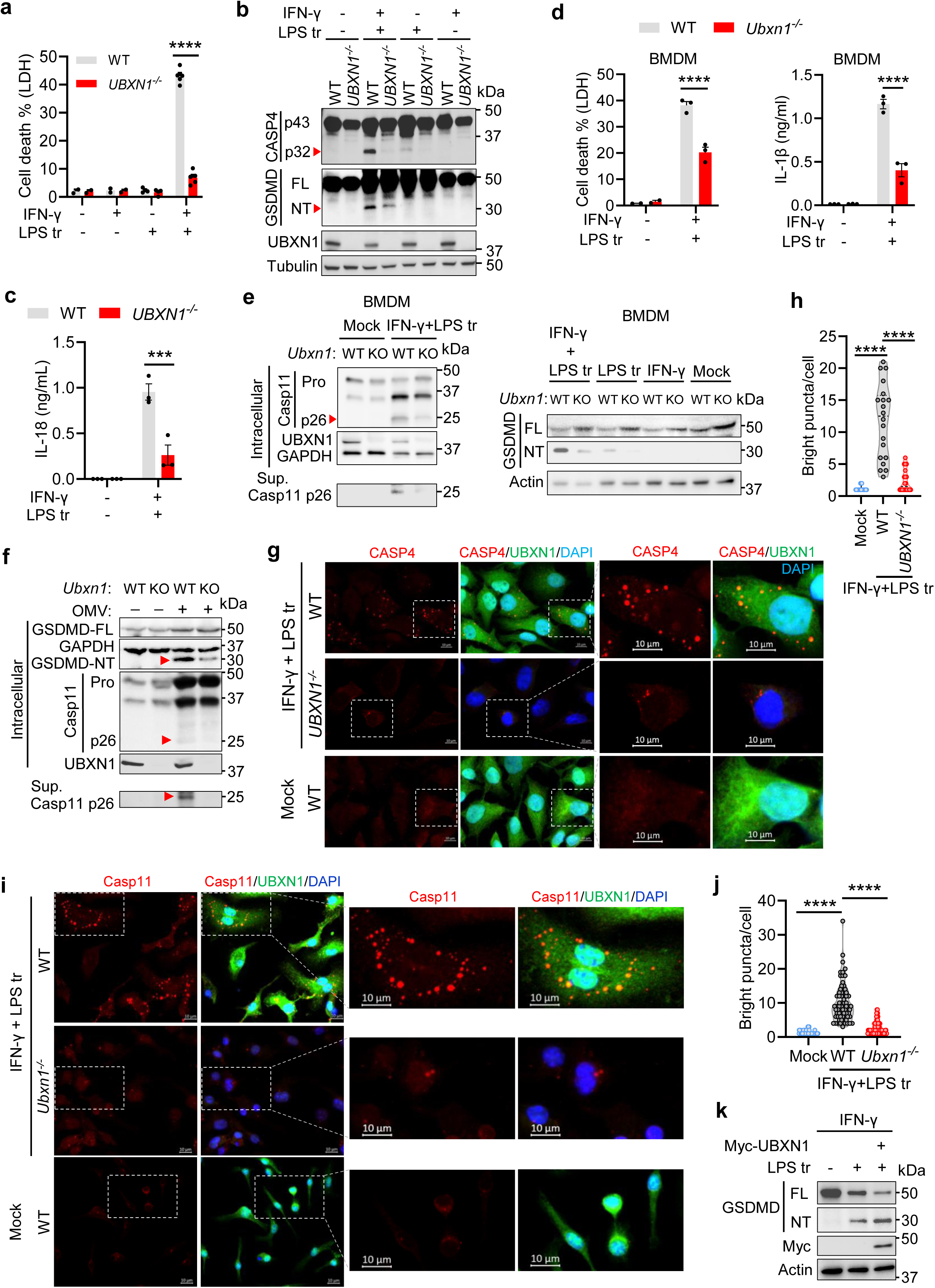
UBXN1 is critical for non-canonical inflammasome signaling. (**a**) Cell death (LDH release) of HeLa cells that were primed with (+) / without (-) 10 ng/mL of human IFN-γ for 12 h and/or transfected with 1 μg/mL LPS (LPS tr) for 5 h. (**b**) Immunoblots of caspase-4 and GSDMD in HeLa treated as in (**a**). The red arrow heads indicate cleaved products. FL, full-length; NT, N-terminal fragment. (**c**) The concentrations of IL-18 in the culture medium of HeLa cells treated as in (**a**). (**d**) Cell death and IL-1β released by BMDMs that were primed with murine IFN-γ (10 ng/mL) for 3 h and then transfected with LPS (1 μg/mL) (LPS tr) for 16 h. N=3 mice/group. (**e**) Immunoblots of caspase-11 and GSDMD in BMDMs treated as in (**d**). Sup: supernatant with secreted p26. (**f**) Immunoblots of caspase-11 and GSDMD in BMDMs treated with outer membrane vesicles (OMV) derived from *E. coli* for 12 h. The red arrow heads indicate desired bands. (**g**) Immunofluorescent confocal images of caspase-4, UBXN1 and DNA (stained with DAPI) in HeLa cells treated as in (**a**). A region of interest (ROI) from each image is enlarged. Mock: transfection reagent only. Scale bar: 10 µm. (**h**) The violin plot of bright caspase-4 puncta per cell from (**g**). N=20 cells per group, each dot represents one cell. (**i**) Immunofluorescent confocal images of caspase-11, UBXN1 and DNA in BMDMs treated as in (**d**). A ROI from each image is enlarged. Scale bar: 10 µm. (**j**) The violin plot of bright caspase-11 puncta per cell from (**i**). N=70 cells per group, each dot represents one cell. (**k**) Immunoblots of GSDMD in HeLa cells transfected with or without Myc-UBXN1 for 24 h, and treated with human IFN-γ and/or LPS as in (**a**). Data are presented in mean ± S.E.M, each symbol represents a biological replicate in (**a**, **c**, **d**), \*\*\**p*<0.001, \*\*\*\**p*<0.0001 by two-tailed unpaired Student’s *t*-test (**a**, **c**, **d**) or One-way ANOVA (**h**, **j**).

Next, we wanted to recapitulate the results from HeLa cells using primary BMDMs. Because of potential embryonic lethality of the *Ubxn1* null mutation (*26*), a tamoxifen-inducible global knockout mouse model was generated by crossing *Ubxn1^flox^*^/flox^ (flanking exons 3-9) with a Cre recombinase-estrogen receptor T2 (ERT2) line (Supplemental **Fig.S3a**, **b**). To minimize heterogeneity among individual mice, we pooled BMDMs from three ERT2-Cre^+^ *Ubxn1^flox^*^/flox^ mice, treated half of the cells with 4-hydroxyl (OH) tamoxifen to induce *Ubxn1* gene deletion (*Ubxn1*^-/-^) and the other half with the solvent (dimethyl sulfoxide, DMSO) (WT) *in vitro* (Supplemental **Fig.S3c**). Indeed, similarly with HeLa cells, cell death, IL-1β secretion, activation of caspase-11 (both intracellular and secreted p26) and GSDMD (N-terminal fragment) were reduced in *Ubxn1*^-/-^ compared to WT BMDMs following LPS transfection (**Fig.2d, e**). The same result was observed when the caspase-11 inflammasome was activated by *E. coli*-derived outer membrane vesicles (OMVs) (**Fig.2f**). Moreover, by confocal microscopy we observed far fewer caspase-4/11 puncta in UBXN1 knockout than WT HeLa cells and BMDMs, respectively (**Fig.2g-j**; Supplemental **Fig.S3d**). Lastly, complementary to the results of UBXN1 knockout, overexpression of UBXN1 enhanced GSDMD cleavage following LPS transfection (**Fig.2k**). In sum, these results show that UBXN1 positively regulates the non-canonical inflammasome in human and murine cells.

### UBXN1 is dispensable for IFN-**γ**, TLR4 and NLPR3 inflammasome signaling

To trigger robust non-canonical inflammasome signaling, BMDMs and HeLa cells were primed with IFN-γ followed by transfecting LPS. IFN-γ signaling induces caspase-4/11 and guanylate binding protein (GBPs) etc., while LPS stimulates TLR4 signaling that can also primes inflammasomes. Therefore, we asked if UBXN1 targets the IFN-γ receptor-JAK1 (Janus kinase 1)-STAT1 (signal transducer and activator of transcription) or TLR4 signaling pathways. We observed no significant difference in JAK1 and STAT1 phosphorylation, *Tnf* or *Casp11* mRNA levels in IFN-γ-primed WT and *Ubxn1*^-/-^ BMDMs (Supplemental **Fig.S4a**, **b**). Moreover, human IFN-γ-induced GBP expression was largely comparable between WT and *UBXN1*^-/-^ HeLa cells (Supplemental **Fig.S4c**, **d**). These observations demonstrate that UBXN1 is not involved in IFN-γ signaling. LPS-induced *Tnf* mRNA expression was slightly enhanced in *Ubxn1*^-/-^ BMDMs (Supplemental **Fig.S4e**). Overall, these results suggest that UBXN1 directly regulates the non-canonical inflammasome pathway and not through the feed-forward TLR4 signaling pathway.

Next, we induced the NLRP3-caspase-1 inflammasome with LPS+ATP or nigericin in primary BMDMs. Secretion of IL-1β and IL-18 and caspase-1 activation (p10) were similar between WT and *Ubxn1*^-/-^ cells (Supplemental **Fig.S5a**, **b**), and LPS-induced NLRP3 expression was also similar (Supplemental **Fig.S5c**). Hence, UBXN1 is dispensable for the NLRP3 inflammasome signaling.

### UBXN1 facilitates the pathogenesis of LPS and polymicrobial sepsis in mice

The aforementioned *in vitro* results clearly demonstrate an important role for UBXN1 in the non-canonical inflammasome, but its role *in vivo* was unknown and thus we characterized UBXN1 function during caspase-11-mediated sepsis (*5–7*). After five doses of tamoxifen (Tmx) in ERT2-Cre^+^*Ubxn1*^flox/flox^ mice, UBXN1 protein was effectively depleted in most tissues except the brain (Supplemental **Fig.S6a**). This is likely due to inefficient penetration of Tmx across the blood brain barrier. Nonetheless, this would not impact our study on sepsis. *Ubxn1*^-/-^ (ERT2-Cre^+^*Ubxn1*^flox/flox^ + Tmx) mice showed no discernible, developmental or behavioral abnormalities, when compared to WT or *Ubxn1*^+/+^ (ERT2-Cre^+^*Ubxn1*^flox/flox^ + oil, *Ubxn1*^flox/flox^ + Tmx, *Ubxn1*^flox/flox^ + oil). The major blood immune populations were the same between *Ubxn1*^-/-^ and WT mice (Supplemental **Fig.S6b, c**). Importantly, *Ubxn1*^-/-^ animals were better protected from lethal LPS sepsis than WT or *Ubxn1*^+/+^ littermates (**Fig.3a**, Supplemental **Fig.S7a**). There was no difference in mortality between *Ubxn1^flox/flox^* +Tmx and *Ubxn1^flox/flox^* + oil mice (**Fig.3a**), excluding any side effect of tamoxifen (which was cleared before LPS challenge) on LPS sepsis. Therefore, we used only one WT littermate control, *i.e*., ERT2-Cre^+^*Ubxn1*^flox/flox^ + oil in subsequent experiments. Both WT and *Ubxn1*^-/-^ mice lost ∼18% of body weight by 48 h after LPS, however, the knockout animals recovered faster than WT mice (**Fig.3b**). The body temperature of *Ubxn1*^-/-^ mice dropped more slowly than that of WT mice after LPS (**Fig.3c**). At 36 h after LPS, WT mice tended to huddle together at cage corners, while the knockout animals were more active and agile (Supplemental **Fig.S7b and video**). These results demonstrate that UBXN1 deficiency protects mice from LPS sepsis.

**Fig. 3.**
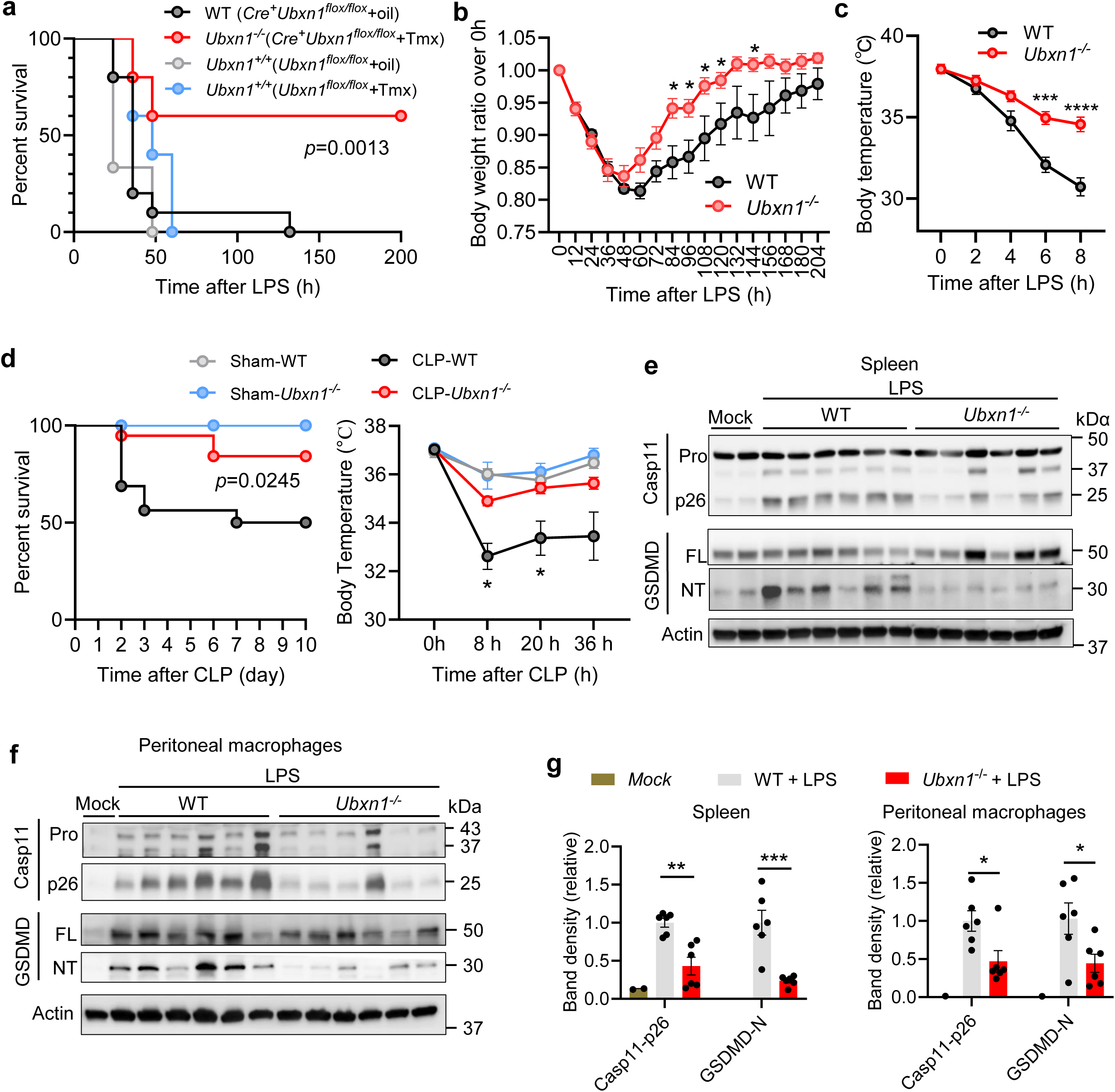
*Ubxn1^-/-^* mice are resistant to lethal LPS and CLP sepsis. (**a**) Survival curves of age- and sex-matched littermates injected with LPS (12 mg/kg) intraperitoneally. N=6 mice for *Ubxn1^flox/flox^* +oil, 5 for *Ubxn1^flox/flox^* +tamoxifen (Tmx), 10 for WT (Cre^+^*Ubxn1^+/+^*+oil) and *Ubxn1^-/-^* (Cre^+^*Ubxn1^+/+^*+Tmx) respectively. *p*=0.0013 (Log-Rank test, WT vs. *Ubxn1^-/-^*). (**b**) Body mass change of WT and *Ubxn1^-/-^* littermates treated with 8 mg/kg of LPS. N=10 mice/genotype. (**c**) Body temperature of WT and *Ubxn1^-/-^* littermates treated with 10 mg/kg of LPS. N=8 mice per group. (**d**) Survival curves and body temperature of age- and sex-matched littermates subjected to cecal ligation and puncture (CLP) sepsis. In the survival curves, N=16 mice for WT and 19 for *Ubxn1^-/-^*, *p*=0.0245 (Log-Rank test). In the temperature curves, N=4 mice for Sham/group and 6 for CLP/group. (**e, f**) Immunoblots of caspase-11 and GSDMD in the spleen and peritoneal macrophages of age- and sex-matched littermates treated with 10 mg/kg of LPS intraperitoneally for 36 h. Each lane represents one mouse. Mock, mice treated with phosphate-buffered saline; Pro, inactive full-lengh; FL, full-length; NT, active N-terminal fragment. (**g**) Quantification of the protein band density in (**e**, **f**). Y-axis shows the ratio of normalized p26 or GSDMD-NT bands to those in group of WT+LPS. Actin was used for internal normalization. N=6 for each LPS-treated group and 2 for Mock. Data are presented in mean ± S.E.M, \**p*<0.05, \*\*\**p*<0.001, \*\*\*\**p*<0.0001 by multiple unpaired Student’s *t*-test in (**b**, **c**, **g**); in the right panel of (**d**), \**p*<0.05, two way-ANOVA, Dunnett comparisons.

In addition to the LPS endotoxemia model, the cecal ligation-and-puncture (CLP) induces polymicrobial, inflammasome-dependent lethal sepsis in mice (*8*) that most closely represents the progression and characteristics of human sepsis (*27*). Consistently, the *Ubxn1*^-/-^ animals had better survival after induction of CLP sepsis than did the WT littermates. The body temperature of *Ubxn1*^-/-^ mice with CLP was also higher than that of WT mice with CLP. As negative controls, all the sham mice survived (**Fig.3d**).

Considering the crucial role of the noncanonical inflammasome in both LPS and CLP sepsis pathogenesis (*5–8*), we investigated caspase-11 and GSDMD activation in mice. Indeed, the levels of active p26 fragment of caspase-11 and N-terminal fragment of GSDMD were lower in the spleen and peritoneal macrophages of *Ubxn1*^-/-^ mice than those in WT following LPS administration (**Fig.3e-g**). Consequently, the concentrations of inflammasome-dependent IL-1β and IL-18 were reduced in the sera and peritoneal lavages of *Ubxn1*^-/-^ mice at various timepoints after LPS (**Fig.4a, b**). The secondary inflammatory mediators that can be induced by IL-1β and IL-18 were also reduced, including IL-6, IL-12, G-CSF, TNF-α etc. (*28–30*). However, IFN-β induction was comparable between WT and knockout mice, further suggesting the dispensable role of UBXN1 in TLR4 signaling (**Fig.4a, b**; Supplemental **Fig.S8**). Similar results were noted in the animals with CLP sepsis (**Fig.4c**; Supplemental **Fig.S8**). In sum, these animal model studies show an important role of UBXN1 in sepsis pathogenesis.

**Fig. 4.**
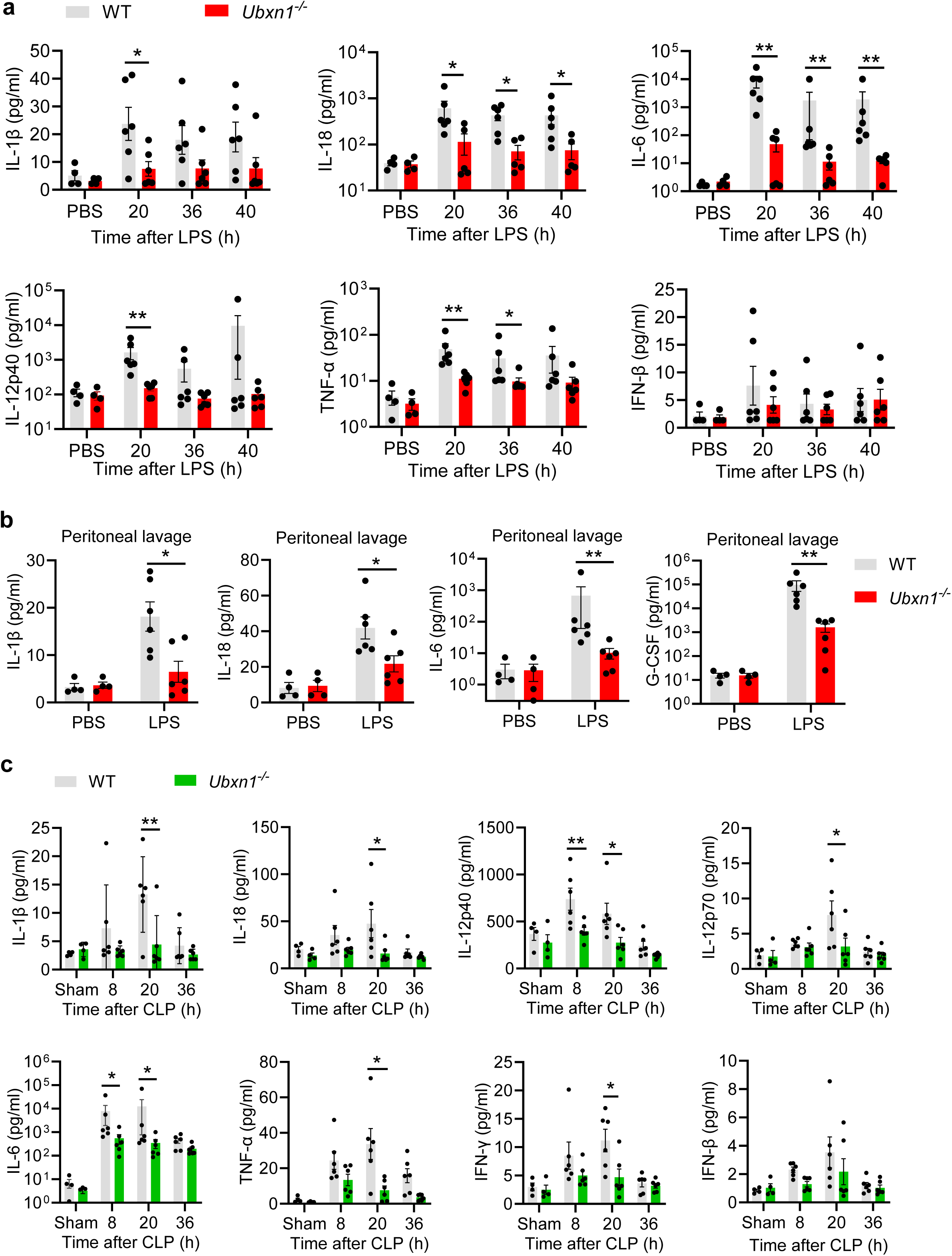
*Ubxn1^-/-^* mice present reduced inflammatory responses during sepsis. (**a**, **b**) The concentrations of cytokines/growth factors in the (**a**) sera at various timepoints and (**b**) peritoneal lavages at 40 h in age- and sex-matched littermates challenged with 10 mg/kg of LPS intraperitoneally. PBS: mice administered with phosphate-buffered saline. (**c**) The concentrations of serum cytokines in age- and sex-matched littermates subjected to sham and a cecal ligation-and-puncture (CLP) procedure. Data are presented in mean ± S.E.M, each dot represents one animal, \**p*<0.05, \*\**p*<0.01 by two way-ANOVA, Dunnett comparisons.

### UBXN1 participates in the caspase-4/11 inflammasome assembly

The aforementioned data suggest that UBXN1 directly regulates the caspase-4/11 inflammasome. Therefore, it was reasoned that endogenous UBXN1 would be associated with the LPS-caspase-4 complex if UBXN1 interacted with caspase-4/11. To address this, we transfected HeLa cells with either biotin or biotin-conjugated LPS followed by streptavidin beads precipitation. Indeed, both UBXN1 and caspase-4 were pulled down together with biotin-LPS (**Fig.5a, b**), aligning with our unbiased mass spectrometry screen for caspase-4 binding partners (**Fig. 1a**). As a negative control, UBXN6, which shares similar domains with UBXN1, was not precipitated by biotin-LPS (**Fig.5b**). Of note, the amount of caspase-4 bound by biotin-LPS was significantly lower in *UBXN1*^-/-^ than WT cells (**Fig.5a**). Consistently, in WT primary BMDMs, UBXN1 and caspase-11 co-localized extensively with FITC-LPS, while the caspase-11 puncta and caspase-11-LPS complexes were significantly reduced in *Ubxn1*^-/-^ cells (**Fig.5c**). These results suggest that UBXN1 participates in the assembly of the caspase-4/11 inflammasome complex. To address this finding more directly, we transiently overexpressed FLAG-CASP4 or FLAG-GSDMD in WT and *UBXN1*^-/-^ HeLa cells to induce pyroptosis without LPS (*31*). The caspase-4-induced pyroptosis was appreciably inhibited in *UBXN1*^-/-^ cells, compared to that in WT cells. However, the GSDMD-induced cell death was the same between the two cell lines (**Fig.5d**), arguing that UBXN1 targets the assembly of the caspase-4 complex and not GSDMD pores. Consistently, compared to the vector (Lane 2 from left), overexpression of FLAG-CASP4 alone activated GSDMD cleavage slightly (Lane 3), and this effect was enhanced in the presence of Myc-UBXN1 (Lane 4) (**Fig.5e**). These results confirm that UBXN1 directly targets caspase-4/11 inflammasome activation.

**Fig. 5.**
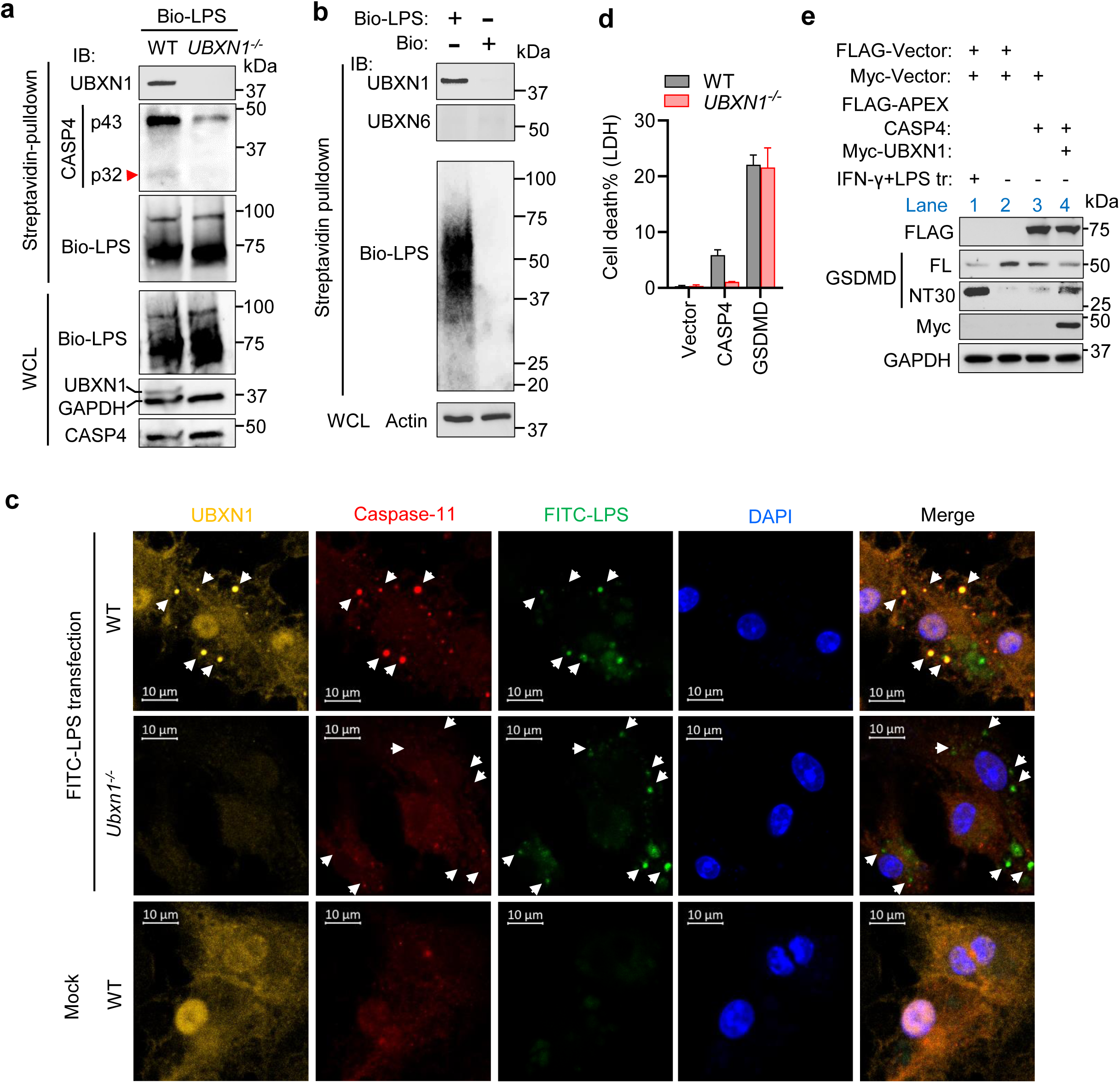
UBXN1 binds and facilitates the caspase-4/11 inflammasome assembly. (**a, b**) Streptavidin pulldown of biotinylated (Bio) LPS from HeLa cells that were primed with IFN-γ and transfected with biotin or Bio-LPS for 5 h. Bio-LPS was detected with a streptavidin-conjugated HRP (horse radish peroxidase). IB, immunoblot. (**c**) Immunofluorescent confocal images in BMDMs that were primed with IFN-γ and transfected with FITC-LPS. DNA was stained by DAPI. Arrows point to colocalizations of FITC-LPS, UBXN1 and caspase-11 in WT, and localization of FITC-LPS in *UBXN1^-/-^* BMDMs. Mock: transfection reagent only. Scale bar: 10 µm. (**d**) The cell death (LDH release) of HeLa cells that were primed with 10 ng/mL of IFN-γ for 12 h and transfected with a CASP4 or GSDMD plasmid for 24 h. Error bar, mean ± S.E.M., N=2 independent experiments. (**e**) Immunoblots of GSDMD in HeLa cells transfected with a FLAG-APEX-CASP4 and/or Myc-UBXN1 plasmid. The cells primed with IFN-γ and transfected with LPS serve as a positive control for GSDMD cleavage. FL, full-length; NT, active N-terminal fragment.

### UBXN1 promotes unanchored K48/63-Ub binding to, and activation of caspase-4/11

Several UBXNs including UBXN1 and UBXN3B are putative adaptor proteins that bridge E3 ubiquitin ligases and their substrates (*32*), interact with polyUb chains of a ubiquitinated (covalently modified) protein (*33*) or unanchored polyUb chains (*34*) through protein-protein interaction. Therefore, we asked if UBXN1 regulates caspase-4/11 function via ubiquitination. We first overexpressed FLAG-APEX-CASP4 and HA-Ub (WT) in HeLa cells and performed immunoprecipitation with an anti-FLAG antibody. Overexpression of HA-Ub enhanced polyUb modification of caspase-4 in both WT and *UBXN1*^-/-^ cells (Lane 4 and 5 vs 3). Of note, LPS transfection further increased polyUb attachment to caspase-4 in WT (Lane 6), but not in *UBXN1*^-/-^ cells (Lane 7) (**Fig.6a**), suggesting that LPS-elicited polyUb attachment to caspase-4 is dependent on UBXN1. Consistent with this, overexpression of UBXN1 induced obvious polyUb attachment to caspase-4 even without LPS transfection (Lane 4 and 6 vs 3); the ubiquitination was further enhanced by LPS (Lane 10 and 12 vs 4 and 6) (**Fig.6b**). However, overexpression of UBXN3B did not (Lane 5 and 11), suggesting that polyUb conjugation to caspase-4 is unique to UBXN1. To pinpoint the specific linkage of Ub attachment to caspase-4, we overexpressed HA-tagged WT or individual mutant Kn-Ub, FLAG-APEX-CASP4, and Myc-UBXN1 (or vector) simultaneously. Each Kn mutant contains only one active K with the remaining K mutated to R (Supplemental **Fig.S9a**). UBXN1 overexpression enhanced only K48- (Lane 12 vs 13 in the IP panel), K63- (Lane 16 vs 17) as well as WT-Ub of caspase-4 (Lane 2 vs 3) when compared to their corresponding vector controls (**Fig.6c**). Next, testing of endogenous K48/63-Ub attachments to caspase-4 using Ub linkage specific antibodies confirmed that UBXN1 also promoted K48/63-Ub attachment to caspase-11, but not caspase-1 (**Fig.6d**, Supplemental **Fig.S9b**, **c**).

**Fig. 6.**
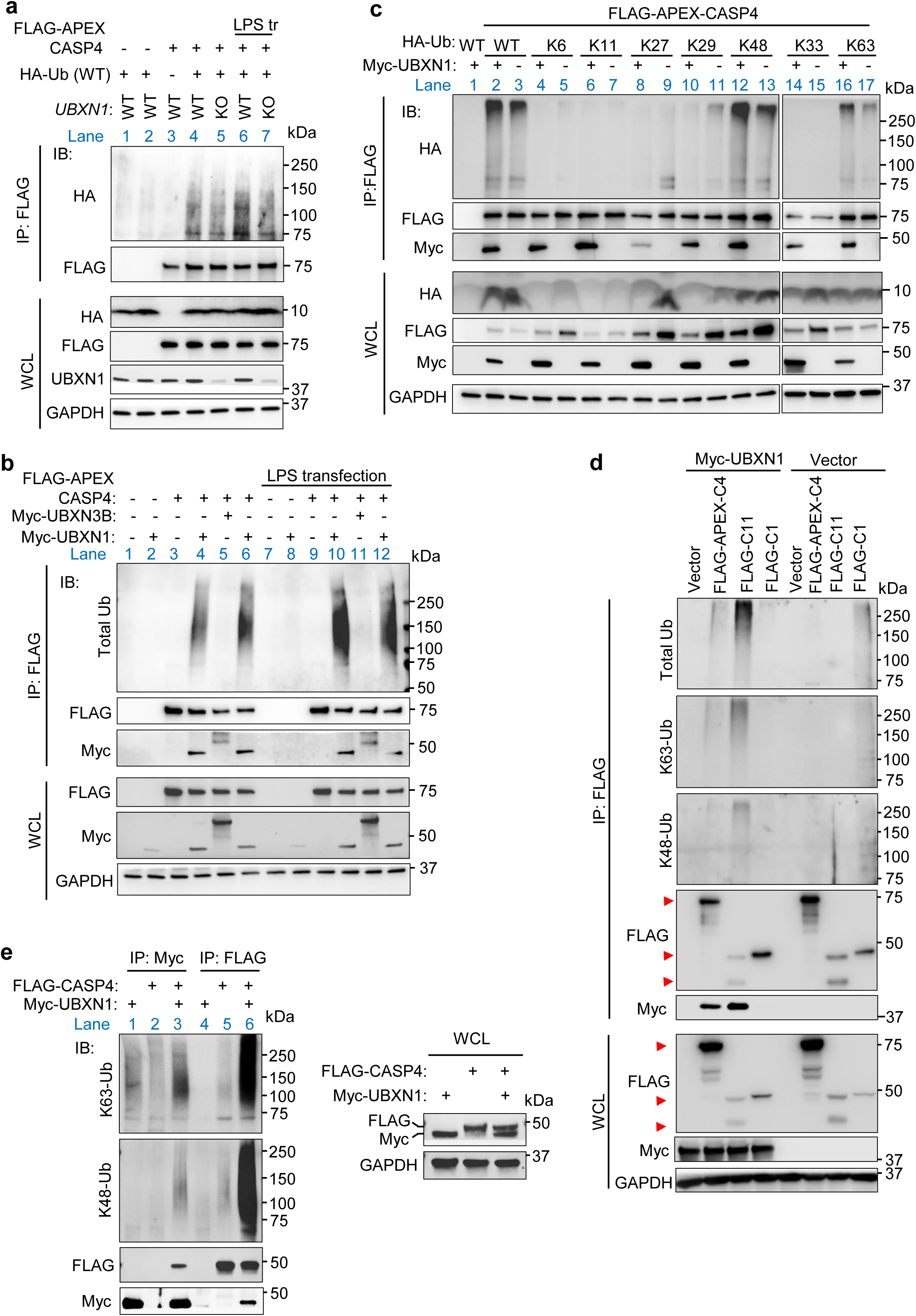
UBXN1 promotes K48- and K63-linked polyubiquitin attachment to caspase-4/11. (**a**) Immunoprecipitation (IP) of FLAG-APEX-caspase-4 with an anti-FLAG antibody. HeLa cells were transfected with a FLAG-APEX-CASP4 and/or HA-Ub (WT) plasmid for 24 h, primed with IFN-γ for 12 h and/or transfected with LPS for 5 h. (-) stands for an empty vector plasmid. Shown are the immunoblots (IB) of indicated proteins. WCL, whole cell lysate; KO, knockout of *UBXN1*. (**b**) IP of FLAG-APEX-caspase-4 with an anti-FLAG antibody from HeLa cells transfected with different combinations of FLAG-APEX-CASP4, Myc-UBXN1, Myc-UBXN3B and corresponding vector (-) for 24 h, followed by treatment with (+) / without (-) human IFN-γ and LPS as in (**a**). The endogenous Ub were examined by polyubiquitin antibodies. (**c**) IP of FLAG-APEX-caspase-4 with an anti-FLAG antibody from HeLa cells transfected with various combinations of FLAG-APEX-CASP4, Myc-UBXN1, HA-tagged WT or individual Kn-Ub (mutant) plasmids. (-) stands for an empty vector plasmid. (**d**) IP of FLAG-caspases with an anti-FLAG antibody from HeLa cells transfected with a FLAG-CASP, Myc-UBXN1 or empty vector plasmid. The red arrow heads indicate correct bands; caspase-11 shows in two bands. The endogenous total, K48, and K63 Ub were detected by specific antibodies. (**e**) IP of FLAG-caspase-4 and Myc-UBXN1 from HEK293T cells. Cells were transfected with the FLAG-CASP4, Myc-UBXN1 or both for 24 h; the cell lysates were equally split for IP with an anti-Myc and anti-FLAG antibody separately. The endogenous K48- and K63-Ub were detected by Ub linkage specific antibodies. Immunoblots (IB) in (**a**-**e**) shows the indicated proteins detected with specific antibodies. WCL, whole cell lysate.

When co-overexpressed, UBXN1 bound caspase-4 at approximately a 2:1 molar ratio (**Fig.1e**); therefore, we questioned if UBXN1 *per se* was ubiquitinated and pulled down by caspase-4. We expressed Myc-UBXN1, FLAG-CASP4 individually or together, and performed immunoprecipitation with an anti-Myc or FLAG antibody. In Myc immunoprecipitation, FLAG-caspase-4 alone was a negative control. As shown in **Fig.6e**, the amount of endogenous K48/63-Ub associated with Myc-UBXN1 alone (Lane 1) was marginally higher than FLAG-CASP4 alone (Lane 2). Notably, much more K48/63-Ub was associated with Myc-UBXN1 in the presence of FLAG-caspase-4 (Lane 3). Conversely, in FLAG immunoprecipitation, Myc-UBXN1 alone served as a negative control, and again co-expression of Myc-UBXN1 significantly increased K48/63-Ub attachment to FLAG-caspase-4 (Lane 6 vs 5) (**Fig.6e**). Consistent with the above results, we identified ubiquitin as a potential new caspase-4 interactor by unbiased mass spectrometry (**Fig.1a**). Of note, ubiquitin was significantly enhanced by UBXN1 overexpression among all the putative caspase-4 interactors involved in ubiquitination (**Fig.7a, b**; Supplemental **Fig.S10a**, **b**).

**Fig. 7.**
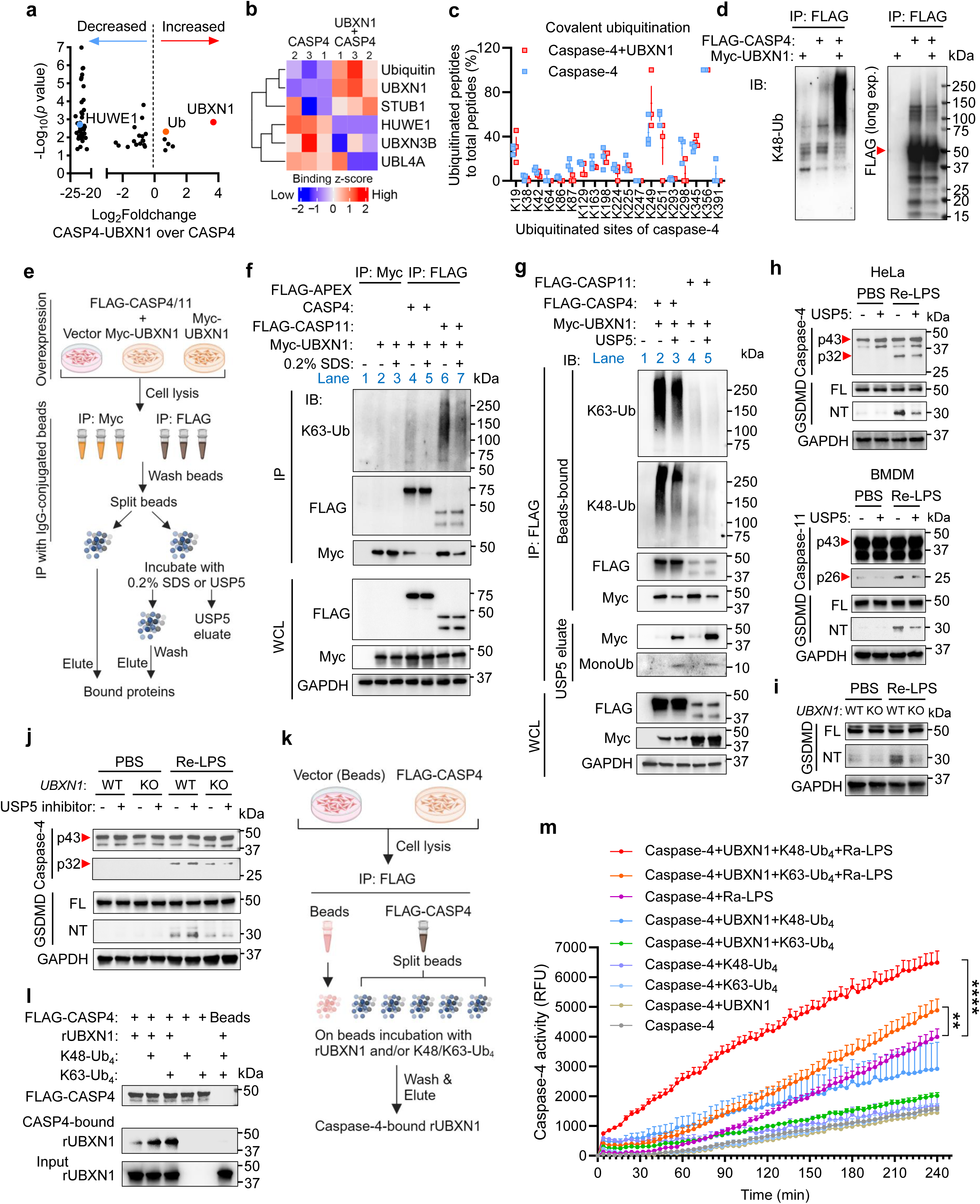
Unanchored K48/63 ubiquitin chains bind and activate caspase-4/11 in a UBXN1-dependent manner. (**a**) A scatter plot of caspase-4 interactors with altered affinity for caspase-4 in the presence of UBXN1. HEK293T cells were primed with IFN-γ for 12 h, transfected with a FLAG-CASP4 and vector or Myc-UBXN1 plasmid for 24 h. Immunoprecipitation (IP) was performed with an anti-FLAG antibody and bound proteins were identified by mass spectrometry. A Log_2_foldchange (FC) represents the ratio of average precursor intensity of a caspase-4 bound protein in the FLAG-CASP4 + Myc-UBXN1 to FLAG-CASP4 + vector group. Proteins involved in ubiquitination are labeled. Dots represent mean of 3 independent experiments, *p*<L0.05 by two-tailed unpaired Student’s t-test with Benjamini–Hochberg. (**b**) The heatmap of binding Z-scores of caspase-4-binding, ubiquitination-related proteins from (**a**). (**c**) The frequency of covalent ubiquitination of individual caspase-4 lysine (K) residues. IP and MS were performed exactly as in (**a**). The ubiquitination ratio of a K is expressed as percent (%) of modified peptide (di-glycine residue) over total peptide counts containing the corresponding K. Error bar, mean ± S.E.M., N=3 independent experiments, no significance by two-way ANOVA, Bonferroni post-hoc test. (**d**) IP of FLAG-caspase-4 from HEK293T cells treated as in (**a**). Samples were immunoblotted (IB) with an anti-K48 Ub linkage specific antibody (left) and anti-FLAG M2 antibody (right). The arrowhead indicates unmodified FLAG-caspase-4. (**e**) The workflow to distinguish free polyubiquitin chain binding to caspase-4/11 from covalent ubiquitination of caspase-4/11. USP5: recombinant ubiquitin-specific proteinase 5 protein. Created with BioRender.com. (**f, g**) The IB of indicated proteins from (**e**), WCL, whole cell lysate. (**h**) Immunoblots of caspase-4/11 and GSDMD from HeLa and primary BMDMs. Cells were primed with IFN-γ, lysed, treated with Re-LPS in the presence (+) or absence (-) of a recombinant USP5 protein for 4 h. PBS, phosphate-buffered saline; FL, full-length; NT, cleaved N-terminal fragment. (**i**) Immunoblot of GSDMD in HeLa cells treated as in (**h**) without USP5. KO, knockout of *UBXN1*. (**j**) Immunoblots of caspase-4 and GSDMD in HeLa cells treated as in (**h**) except that the recombinant USP5 was replaced by an USP5 inhibitor. (**k**) The schematic workflow for *in vitro* protein-protein interaction. Created with BioRender.com. K48-, K63-Ub_4_ are *E. coli*-derived recombinant tetra ubiquitin chains. Recombinant 6xHis-UBXN1 (rUBXN1) is *E. coli-*derived too. (**l**) The immunoblots of indicated proteins in the eluates from (**k**). (**m**) *In vitro* caspase-4 activity assay with a fluorogenic substate, purified insect-derived caspase-4, *E. coli*-derived 6xHis-UBXN1 and Ub_4_, and Ra-LPS. Data are presented in mean + S.E.M., N=3 biological replicates, \*\**p*<0.01, \*\*\*\**p*<0.0001 by repeated measures two way-ANOVA, Sidak’s multiple comparisons test.

Traditionally, ubiquitination is regarded as the covalent modification of a protein by ubiquitin. However, cells also have unanchored polyUb chains that can noncovalently bind and regulate the function of target proteins (*35*). Therefore, we first asked if UBXN1 overexpression promoted covalent ubiquitination of caspase-4 and further attempted to identify its ubiquitination site by immunoprecipitation and mass spectrometry. We observed 19 ubiquitinated lysine residues in FLAG-caspase-4, however, none of them were significantly enhanced when Myc-UBXN1 was co-overexpressed (**Fig.7c**), suggesting that UBXN1 promotes non-covalent ubiquitination of caspase-4. Consistent with this, when detected with an anti-FLAG antibody, the density of the bands above FLAG-caspase-4 (putative ubiquitinated form) relative to FLAG-caspase-4 was the same with/without Myc-UBXN1 (right panel, **Fig.7d**). However, much more K48-Ub was observed by its specific antibody in the presence of Myc-UBXN1 (left panel, **Fig.7d**). These results strongly support the idea that UBXN1 chaperones free polyUb chains to attach to caspase-4. To further confirm this, 0.2% sodium dodecyl-sulfate (SDS) was included in the immunoprecipitation assay to disrupt regular protein-protein interactions, but not the strong antibody-antigen interaction (**Fig.7e**). Indeed, 0.2% SDS did not affect immunoprecipitation of FLAG-caspase-4/11 with an anti-FLAG or Myc-UBXN1 with an anti-Myc antibody (IP, Lane 2 vs 3, 4 vs 5, 6 vs 7), but strongly reduced the amount of Myc-UBXN1 and K63-Ub co-precipitated with FLAG-caspase-4/11 (Lane 4 vs 5, and 6 vs 7) (**Fig.7f**). It was also noted that there was much more free K63-Ub bound to FLAG-caspase-11 than FLAG-APEX-caspase-4, relative to their IP input (Lane 4 vs 6) (**Fig.7f**). This difference could be due to the large APEX tag (27 kDα) that may interfere with the interaction between free polyUb and caspase-4. Indeed, relative to their IP input, FLAG-caspase-4 pulled down the similar amount of K48/63-Ub as FLAG-caspase-11 did (IP, Lane 2 vs 4) (**Fig.7g**). Next, we used USP5 (also known as isopeptidase T, IsoT) to specifically digest unanchored polyUb chains without disrupting regular protein-protein interactions (**Fig.7e**). USP5 obviously reduced the amount of free K48/63-Ub bound to FLAG-caspase-4/11 (Beads-bound, Lane 2 vs 3, and 4 vs 5). This reduction was correlated with the increase of mono ubiquitin in the USP5 eluate (**Fig.7g**). Notably, the binding of Myc-UBXN1 to FLAG-caspase-4/11 was also disrupted by USP5. These results demonstrate that UBXN1 mediates unanchored K48/63-Ub binding to caspase-4/11 and stable caspase-4/11 binding to UBXN1 requires unanchored K48/63-Ub.

### Unanchored K48/63-Ub promote caspase-4/11 activation

Next, we asked if unanchored polyUb chains play a role in caspase-4/11 activation. Because unanchored polyUb can be derived from *de novo* synthesis by specialized E2/E3 and deubiquitylation of substrates by numerous DUBs (*35*), it is challenging to genetically deplete intracellular unanchored polyUb. Our knowledge of unanchored polyUb has been largely obtained from research on USP5 (IsoT) (*35*); however, genetic manipulation of USP5 can alter many cellular activities due to impaired proteasomal protein degradation. To overcome this, we developed a cell-lysate system for the caspase-4/11 inflammasome. We primed HeLa with IFN- γ, lysed cells by douncing, and removed nuclei, large membranes and debris by centrifuging at 14,000 x g. The cleared lysate was then incubated with WT or mutant LPS of various concentrations (Supplemental **Fig.S11a**, **b**). We noted that the Rd and Re mutants without the outer core and O-antigen more potently activated caspase-4 and GSDMD than WT and the Ra mutant (Supplemental **Fig.S11c**, **d**). Using this cell-lysate system, we then investigated if USP5 inhibits the caspase-4 inflammasome by depleting free polyUb chains. On one hand, pretreating the WT cell lysate with a recombinant USP5 protein resulted in a notable reduction in Re-LPS - activated caspase-4/11 and GSDMD cleavage (**Fig.7h**). On the other hand, inhibiting endogenous USP5 enhanced inflammasome signaling (**Fig.7j**). However, the effect of either recombinant USP5 or the USP5 inhibitor was lost in *UBXN1*^-/-^cells as compared to WT controls (**Fig.7i, j**; Supplemental **Fig.S11e**), suggesting that free K48/63-Ub require UBXN1 to regulate caspase-4/11.

To strengthen the concept that unanchored K48/63-Ub and UBXN1 directly cooperate to regulate the non-canonical inflammasome, an *in vitro* caspase-4-UBXN1 binding and caspase-4 activation assay was performed. We first expressed and purified FLAG-caspase-4 from HEK293 cells, then incubated it with *E. coli*-derived recombinant UBXN1 and Ub_4_ (**Fig.7k**). Compared to the control (anti-FLAG beads), FLAG-caspase-4 pulldown of recombinant UBXN1 was significantly enhanced in the presence of either K48-Ub_4_ or K63-Ub_4_ (**Fig7l**). Further, the basal activity of recombinant caspase-4 (expressed and purified from insect cells) increased in the presence of both recombinant UBXN1 and Ub_4_, compared to individual UBXN1 or Ub_4_. Ra-LPS-stimulated caspase-4 activity was also enhanced by UBXN1 and Ub_4_ (Red and orange vs purple lines) (**Fig7m**).

## DISCUSSION

Although *in vitro* caspase-4/11 can be directly activated by LPS, in the complex intracellular environment, efficient activation of caspase-4/11 may require additional cellular factors (*36*). Guanylate binding proteins (GBPs) could serve as a platform to assemble free cytosolic LPS and caspase-4/11 (*37*). However, most studies suggest that GBPs assemble on the surface of Gram-negative bacteria into polyvalent signaling platforms for recruiting caspase-4/11 (*38–40*). In this study, we have discovered that UBXN1 and unanchored K48/63-Ub together promote caspase-4/11 inflammasome assembly.

UBXN1 has been shown to inhibit RIG-I like receptor (RLR) signaling (the primary viral RNA sensor) (*41*) and NF-κB activity (*26, 42*); regulate endoplasmic reticulum-associated degradation (ERAD) positively or negatively (*43–49*); promote the formation of aggresomes (*50*) and mitophagy (*51*). However, all these studies were conducted *in cellulo*, lacking the *in vivo* physiological relevance. With an inducible *Ubxn1* knockout mouse model, we have shown that *Ubxn1* deficiency renders animals resistant to caspase-11 inflammasome-dependent LPS sepsis and polymicrobial CLP sepsis (*5–8*), impairs caspase-11 and GSDMD cleavage and secretion of inflammasome-dependent cytokines. The RLR-inhibiting function of UBXN1 has been recapitulated in macrophages too, however, *Ubxn1* knockout confers no protection against lethal infection of an RNA virus (Data not shown). These observations suggest that UBXN1 primarily regulates the noncanonical inflammasome *in vivo*. Nonetheless, the other UBXN1-regulated pathways could influence sepsis pathogenesis too. For instance, mitophagy inhibits inflammatory responses and alleviates tissue damage during sepsis (*52*), while NF-κB activity does the opposite (*53*). However, UBXN1 seems to regulate mitophagy positively (*51*) and NF-κB activity negatively (*26, 42*), indicating that the primary target of UBXN1 is the noncanonical inflammasome during sepsis.

Unanchored polyUb chains are synthesized *de novo* by specialized ubiquitin conjugating enzyme (E2) and ligase (E3) pairs or derived from deubiquitylation of substrates (*35*). However, the physiological functions of unanchored polyUb chains have been only recently appreciated. Earlier, unanchored polyUb chains were believed to be toxic to cells, due to inhibition of proteasomes and subsequent accumulation of undesirable proteins (*54–58*). Recent studies, however, have shown their important functions in immune pathways and stress responses. For instance, unanchored K63-Ub chains bind TAB2 [TAK1 (transforming growth factor-beta activated kinase 1)-binding protein 2], leading to activation of the TAK1, IKK (IκB Kinase) complex and NF-κB (Nuclear factor κB) in the IL-1 receptor signaling cascade (*59*). K63-Ub decorated with Met1-Ub (namely linear polyUb) bind NEMO (IKKγ/NF-κB essential modulator), resulting in formation of the A20-NEMO complex and termination of NF-κB signaling in the Toll-like receptors (TLRs) and TNFR (tumor necrosis factor receptor) pathways, respectively (*60–62*). Unanchored K63-Ub bind RIG-I (retinoic acid-induced gene I)-like receptors (RLRs) to promote immune responses to RNA viruses (*63, 64*). Unanchored K48-Ub bind IKK-ε to activate itself and STAT1 phosphorylation (signal transducer and activator of transcription 1), leading to interferon (IFN) expression and signaling, respectively (*65*). Our study, however, extends the function of free polyUb to inflammasomes. Caspase-4/11 interacts with both free K48-Ub and K63-Ub chains via UBXN1, where UBXN1-caspase-4/11 binding is reliant on unanchored polyUb. This mode of interaction and regulation may minimize spontaneous activation by abundant unanchored polyUb chains *per se* in cells. For example, a cell may contain ∼6000 unanchored K63-Ub molecules at steady state and requires only ∼30 molecules for full activation of RIG-I (*63*). Free K48-Ub chains may be even much more abundant than K63-Ub. Nonetheless, accumulation of unanchored ubiquitin chains in the vicinity of a target may be necessary for efficient binding and activation of the target. This can be achieved by bringing a specialized E3 ligase like TRIM25 (tripartite motif containing 25) or TRAF6 (TNF receptor associated factor 6) to provide *de novo* synthesis of unanchored polyUb to RLRs (*63*) or TAB2 (*59*), respectively. Alternatively, a ubiquitin chain-binding protein like UBXN1 can enrich and stabilize unanchored polyUb chains around a target like caspase-4/11. Indeed, a very recent work showed that UBXN1 strongly bound unanchored K48-Ub_4_, and to a lesser extent, K63-Ub_4_ (*34*). These data may explain why caspase-4/11 is not selective in the Ub linkage type. Nonetheless, free K48-Ub chains seems more potent in activating caspase-4 than K63-Ub (**Fig.7m**). Moreover, cellular free K48-Ub chains may be more abundant than free K63-Ub, thus more important in activating caspase-4/11.

In sum, this study identifies UBXN1 as a positive regulator of the noncanonical inflammasome and establishes unanchored polyUb as the “glue” that complexes the noncanonical inflammasome into its active structure. Most importantly, UBXN1 plays a notable role in polymicrobial CLP sepsis that closely resembles the progression and characteristics of human sepsis (*27*). Indeed, UBXN1 expression was significantly increased in patients experiencing life-threatening sepsis (*66*). Therefore, future work intends to investigate UBXN1 as a potential drug target for human septicemia.

## MATERIALS AND METHODS

### Mice

All the animal procedures were approved by the Institutional Animal Care and Use Committee at UConn Health adhering to federal and state laws. We applied a CRISPR-Cas9 method to generating a LoxP flanked *Ubxn1*^flox/flox^ mouse line at UConn Health. Briefly, a *loxP* sequence (ATAACTTCGTATAATGTATGCTATACGAAGTTAT) was inserted into the intron between Exon 2 and 3 (5’-*loxP*), and the intron between Exon 9 and 10 (3’-*loxP*), the C57BL/6 embryonic stem cell genome. Appropriately modified ES were injected into C57BL/6 mice. The *Ubxn1*^flox/flox^ mouse line was crossed with a tamoxifen inducible Rosa26-CreERT2 line (The Jackson Laboratory, Stock # 008463), to generate *ERT2*-Cre^+^ *Ubxn1*^flox/flox^ mice. Genotyping was performed with genomic DNA and Choice Taq Blue MasterMix (Denville Scientific, Cat# CB4065-8) under the following PCR: 95□°C for 1 s, 34 cycles of 94□°C for 1□min, 60□°C for 30□s, 72□°C for 30□s, and then 72□°C for 7□min, 4□°C to stop. The primers for genotyping 5’- loxp were F: 5’-ATCGAGATGGGCTTTCCCAG-3’ and R: 5’-GTAGGCAGGGGTTGGTTAGC-3’. The PCR products were 303 bp (mutant allele) and 267 bp (WT allele). The primers for genotyping 3’-loxp were F: 5’-CTGTGCAGTTGCTCAGTGG-3’ and R: 5’- TCAGCTGGGACACTTCTTGG-3’. The PCR products were 330 bp (mutant allele) and 294 bp (WT allele). The Cre primers were common-5’ AAGGGAGCTGCAGTGGAGTA, WT reverse, 5’- CCGAAAATCTGTGGGAAGTC, and mutant reverse, 5’-CGGTTATTCAACTTGCACCA. The PCR resulted in products of 297□bp (WT) and/or 450□bp (Cre). To induce global *Ubxn1* knockout, 1 mg of tamoxifen (dissolved in corn oil) was administered to each mouse (>6 weeks old) every other day for five times in total. *ERT2*-Cre^+^ *Ubxn1*^flox/flox^ mice treated with corn oil served as the wild-type control (*Ubxn1*^+/+^). The mice then rested for at least two weeks to allow for tamoxifen clearance before being employed for examination of knockout efficiency and any further analysis.

### Reagents and antibodies

Antibodies for GAPDH (Cat# 60004-1-Ig, 1:2000), UBXN1 (Cat# 16135-1-AP, 1:1000), UBXN6 (Cat# 14706-1-AP, 1:1000), GBP1 (Cat# 15303-1-AP, 1:1000) were purchased from Proteintech Group. Caspase-4 (Cat# 4450S, 1:1000), Gasdermin D (Cat# 39754S, 1:1000), Cleaved Gasdermin D (Asp275) (Cat# 36425S, 1:500), Cleaved Gasdermin D (Asp276) (Cat# 10137S, 1:1000), NLRP3 (Cat# 15101S, 1:1000), Tubulin (Cat# 2144S, 1:2000), Actin (Cat#4967S, 1:2000), UBXN3B (Cat# 34945S, 1:1000), K63-linkage specific polyubiquitin (Clone D7A11, Cat# 5621S,1:500), K48-linkage specific polyubiquitin (Clone D9D5, Cat# 12805S, 1:1000), Phospho-Jak1 (Tyr1034/1035) (Cat# 74129S, 1:1000), Phospho-Stat1 (Tyr701) (Cat# 9167S, 1:1000), Myc-Tag (Cat# 2276S, 1:2000) and HA-Tag (Clone C29F4, Cat# 3724S, 1:1000) antibodies were from Cell Signaling. Caspase-1 (Cat# ab179515, 1:1000), Caspase-11 (Cat# ab180673, 1:1000), GSDMD (Cat# ab209845, 1:1000) antibodies and recombinant human USP5 protein (Active) (IsoT, Cat# ab269124) were from Abcam. The mouse anti-Myc antibody (Clone 9E10, Cat# TA150121, 1:1000) was obtained from OriGene Technologies. Antibodies for HUWE1 (Cat# A20708, 1:1000), STUB1 (Cat# A11751, 1:1000), K29-linkage specific polyubiquitin (Cat# A18198, 1:500), K33-linkage specific polyubiquitin (Cat#A18199, 1:500) were purchased from ABclonal. The Anti-FLAG M2 magnetic beads (Clone M2, Cat# M8823), Monoclonal Anti-FLAG M2 antibody (Cat# F1804, 1:2000), Lipopolysaccharides (*E. coli* O111:B4, Cat# L3024), LPS-Ra mutant (*E. coli* EH100, Cat# L9641), LPS-Rd mutant (*E. coli* F583, Cat# L6893), LPS-FITC conjugate (*E. coli* O111:B4, Cat# F3665), Biotin (Cat# B4501), Nigericin (Cat# N7143), ATP (Cat# A6419) and Tamoxifen (Cat# T5648) were obtained from Sigma-Aldrich. LPS-Re mutant (Lipid A, *E. coli* R515, Cat# ALX-581-007-L001) was from Enzo. Recombinant INF-γ protein of mouse (Cat# 485-MI-100) and human (Cat# 285-IF-100), Recombinant human INF-β were obtained from R&D systems. The anti-Myc magnetic beads (Clone 9E10, Cat# 88842), Donkey anti-Mouse Alexa Fluor™ Plus 488 (Cat# A32766, 1:200), Donkey anti-rabbit Alexa Fluor™ Plus 594 (Cat# A32754, 1:200), Ubiquitin Polyclonal Antibody (Cat# 1859660, 1:1000), DAPI Solution (Cat# 62248, 1:1000), Pierce™ High Capacity Endotoxin Removal Spin Columns (Cat# 88273), PureLink™ Genomic DNA Mini Kit (Cat# K182002), HiPure Plasmid Filter Midiprep Kit (Cat# K210014), and Quick Plasmid Miniprep Kit (Cat# K210010) were from Thermo Fisher. K48 Tetra-Ubiquitin (Cat# SBB-UP0070) and K63 Tetra-Ubiquitin (Cat# SBB-UP0073) were a product of South Bay Bio. K63-linked Polyubiquitin (Ub n2-7) (Cat# UBI-60-0109-100) was from Cosmo Bio USA. The TransIT-X2 Dynamic Delivery System (Cat# MIR6005) was from Mirus Bio LLC. Biotinylated LPS (*E. coli* 0111:B4, Cat# tlrl-lpsbiot) was a product of InvivoGen. Streptavidin Magnetic Beads (Cat# S1420S) was a product of New England Biolabs.

### Plasmids

The HA-Ub (WT), HA-K6Ub (all the other K are mutated to R), HA-K11Ub, HA-K27Ub, HA-K29Ub, HA-K33Ub, HA-K48Ub,and HA-K63Ub plasmids were kind gifts of Dr. Rongtuan Lin (*67*). The FLAG-APEX-CASP4 and FLAG-CASP1 plasmids were kind gifts of Dr. Vijay A. Rathinam. The FLAG-CASP4 plasmid was a kind gift of Dr. Jianbin Ruan. The FLAG-Casp11 plasmid was from Addgene (Cat# 21145). The Myc-UBXN1 and Myc-UBXN3B plasmids were described previously (*25, 41*).

### Mouse model of sepsis

The sepsis model was induced by intraperitoneal (i.p.) injection of LPS or cecal ligation and puncture (CLP) as described previously (*68, 69*). For CLP model, age- and sex-matched WT and *Ubxn1^-/-^*mice were subjected to 14 hr fasting before the procedure. The mice were anesthetized by injecting ketamine (100 mg/mL*320 µL) /xylazine (100 mg/mL*100 µL) mixture intraperitoneally (i.p.1.2ml/kg bodyweight). The mice were then fixed gently on a circulating warm operating pad. The abdominal skin was disinfected using 70 % ethanol, and a small (∼1 cm) longitudinal skin incision was made left to midline. The cecum was ligated at 1.5 cm from the end and perforated by 5 times of through-and-through puncture near to the ligation using a 21 G needle, avoiding blood vessels, followed by squeezing the ligated cecum to confirm patency and small amount of feces extruded out. Then, the cecum was repositioned into the peritoneal cavity and the abdominal musculature was closed layer by layer. Sham-operated mice were performed using the same surgical procedure but without ligation or puncture of the cecum. Rehydrate with subcutaneously 37 °C saline in a dose of 0.03 mL/g body weight, and buprenorphine (0.05 mg/kg body weight) was injected subcutaneously immediately after surgery. All mice were free to have access to water and food after the procedure. For sepsis induced by LPS, age-and sex-matched mice were injected intraperitoneally (i.p.) with 8, 10 or 12 mg/kg of E. coli O 111:B4 LPS unless otherwise indicated. Mice injected with same volume of PBS were considered as controls. Body weight, rectal temperature and survival were monitored, and samples for sera, peritoneal lavage and tissues were collected at indicated time points.

### Cell culture and isolation of peritoneal or bone marrow-derived macrophages (BMDMs)

HeLa (Adenocarcinoma epithelial cell, Cat# CCL-2), HEK293T cells (human embryonic kidney, Cat# CRL-3216) and L929 cells (mouse fibroblast cells, Cat# CCL-1) were purchased from the American Type Culture Collection (ATCC) (Manassas, VA 20110, USA). HEK293T, HeLa and L929 cells were grown in Dulbecco’s modified Eagle’s medium (DMEM, Corning, Cat#10-013-CV) supplemented with 10% fetal bovine serum (FBS, Gibco™, Cat#A52567-01) and 1% antibiotics/antimycotics (ThermoFisher Scientific, USA). These cell lines are not listed in the database of commonly misidentified cell lines maintained by ICLAC and have not been authenticated in our hands. They are routinely treated with MycoZAP (Lonza, Switzerland) and tested for mycoplasma contamination in our hands.

Bone marrow cells were isolated and differentiated into macrophages (BMDM) using the established methods (*70*). Briefly, bone marrow cells were obtained from mouse hind legs by centrifugation at 8000rpm for 1 min and differentiated in L929-conditioned medium (RPMI 1640, 20% FBS, 30% L929 culture medium, 1x antibiotics/antimycotics) in a 10-cm Petri dish at 37□°C, 5% CO_2_ for 5-7 days, with a change of medium every 2 days. Attached BMDMs were dislodged by incubating in ice-cold PBS for 30 min followed by pipetting and counted for plating in cell culture plates. The BMDMs were seeded and treated in plating medium (RPMI 1640, 10% FBS, 5% L929-conditioned medium, 1x antibiotics / antimycotics).

For isolation of peritoneal macrophages, the mice were injected intraperitoneally with 5 ml of 3% (w/v) Brewer thioglycollate medium. After 3 days, the mice were euthanized and injected 5 ml of ice-cold PBS into the peritoneal cavity. The abdomen was massaged gently for 2 min to dislodge macrophages. The peritoneal fluid was then collected using a sterile syringe with 25 G needle and transferred to a 15 ml tube on ice. The fluid was filtered through a 70 µm cell strainer to remove debris or tissue, and centrifuged at 500 g, 4°C for 5 min. The pelleted cells were then resuspended in complete DMEM with 10% FBS and cultured in plates at the desired density for 2 h at 37□°C, 5% CO_2._ The primarily macrophages adherent were obtained after a gentle wash with PBS to remove non-adherent cells and treated as requirement.

### Generation of gene knockout cell lines using CRISPR-Cas9

Gene knockout of *UBXN1*, *STUB1*, *HUWE1* and *CASP4* in human cell lines by CRISPR-Cas9 were performed based on our previous studies (*70*). Briefly, pre-designed, gene unique guide (g) RNAs (Integrated DNA Technologies, USA) (Sequences in Supplemental **Table 1**) were subcloned into a lentiCRISPR-V2 vector (*71*) and correct insertion was confirmed by DNA sequencing. To generate lentiviral particles, each gRNA vector was transfected into HEK293T cells with the packaging plasmids pCMV-VSV-G (*72*) and psPAX2 (#12259, from the Didier Trono lab via Addgene, USA). A half of the cell culture medium was replaced with fresh medium at 24 h and viral particles were collected at 48-72 h after transfection. The cell culture supernatant (viral particles) was cleared by brief centrifugation at 1000 rpm for 5 min. Target cells at ∼50% confluence in a 6-well plate were transduced with 1 mL of each lentiviral preparation and 8 μg/mL polybrene for 24 h and selected with 1□µg/mL of puromycin in growth media for 5 days. The cells were then cultured in fresh complete DMEM for experiments or frozen for stork. The knockout efficiency in each cell line was examined by immunoblots with specific antibody.

### Flow cytometry

Flow cytometry was performed based on methods in our published study (*24*). Whole blood cells were token from the retro-orbital sinus of anesthetized mice by ketamine/xylazine. The red blood cells were lysed three times with an RBC lysis buffer (BioLegend, Cat. # 420301) if needed. Pelleted cells were suspended in FACS buffer and stained for 30 min at 4 °C with the desired antibody cocktails (BioLegend) of APC-Fire 750-anti CD11b (Cat. # 101261, clone M1/70), Alexa Fluor 700-anti Ly-6G (Cat. # 127621, clone 1A8), PE-Dazzle 594-anti CD3 epsilon (Cat. # 100347, clone 145-2C11), Brilliant Violet 711-anti CD115 (Cat. # 135515, clone AFS98), Zombie UV (Cat. # 423107), PE-Cy7-anti CD45 (Cat. # 103113, clone 30-F11), APC-anti CD19 (Cat. # 152410, clone 1D3/CD19). After staining and washing, the cells were fixed with 4% PFA and loaded on a Becton–Dickinson FACS ARIA II, CyAn advanced digital processor (ADP). Data were analyzed using the FlowJo software. Among CD45+ cells, CD11b+ Ly6G+ cells were classified as neutrophils, Ly6G− CD11b+ CD115+ as monocytes, CD3+ as total T cells, CD19+ as B cells.

### Induction of non-canonical and canonical inflammasome response

For activating non-canonical inflammasome, HeLa and primary BMDMs obtained as described above were cultured in complete DMEM containing 10% FBS and 1% Penicillin-Streptomycin at 37□°C and 5% CO2. For assessing inflammasome and cell death responses, HeLa cells were primed with 10 ng/mL of human IFN-γ for 12 h, and then transfected with 1 μg/mL LPS (*E. coli* O111:B4) (or Bio-LPS, FITC-LPS for immunofluorescence) for 5 h using TransIT-X2 Dynamic Delivery System. BMDMs were primed with murine IFN-γ (10 ng/mL) for 3 h and then transfected with LPS (1 μg/mL) for 16 h. For inflammasome response induction by bacterial outer membrane vesicles (OMVs), BMDMs were stimulated with 10 μg/mL of *E. coli* OMVs (InvivoGen, Cat# tlrl-omv-1) for 16 h. For plasmid overexpression-induced inflammasome response, HeLa cells were treated with or without IFN-γ priming for 12 h and transfected with FLAG-CASP4, FLAG-APEX-CASP4 and/or Myc-UBXN1 plasmids for 24 h. To stimulate canonical NLRP3 inflammasome, primary BMDMs were primed with LPS (1 μg/ml) for 3 h and then stimulated by ATP (5 mM) or Nigericin (10 μg/ml) for 30 min. Cell lysates and supernatants were collected for immunoblots of caspase-4/11 and GSDMD cleavage or assays of cytokines and LDH release.

### Extraction of total RNA and RT-qPCR

Real time quantitative PCR were performed as in previous study (*73*).BMDM cells were collected in 350□µL of lysis buffer (RNApure Tissue & Cell Kit, CoWin Biosciences, USA), containing 1% β-Me. Total cellular RNA was extracted according to the product manual. Reverse transcription of RNA into complementary DNA (cDNA) was performed using the PrimeScript™ RT reagent Kit (TaKaRa Bio, Inc, Cat# RR037A). Quantitative PCR (qPCR) was performed with gene-specific primers and iTaq Universal SYBR Green Supermix (BioRad, Cat# 1725124). Results were calculated using the –ΔΔCt method and a housekeeping gene (ACTB) as an internal control. The qPCR primers for were list in Supplemental **Table 2**.

### Cell death assays and cytokine assays by individual and multiplex ELISA

Cell death was analyzed by measuring LDH release into the cell culture medium with the Cytotoxicity Detection Kit^PLUS^ (LDH) kit (Roche, Cat# 04744926001) according to the manufacturer’s protocol. The release of IL-1β or IL-18 in cell culture supernatants were quantified by a mouse IL-1β DuoSet ELISA kit (R&D, Cat# DY401-05) or a human IL-18 DuoSet ELISA Kit (R&D, Cat# DY318-05). The levels of cytokines or growth factors in mouse sera, peritoneal lavage and culture medium of primary BMDMs were determined by a LEGENDplex™ bead-based multiplex ELISA kit (Biolegend, USA). The procedures were performed fully according to the product manuals. Briefly, 25 µL of samples or standards were mixed with antibody-coated premix microbeads in a filter-bottom microplate and incubated at room temperature for 2 h with vigorous shaking at 500 g. After removal of unbound analytes and two washes, 25 µL of detection antibody were added to each well, and the plate was incubated at room temperature for 1 h with vigorous shaking at 500 g. 25 µL of SA-PE reagent was then added directly to each well, and the plate was incubated at room temperature for 30 min with vigorous shaking at 500 g. After twice washes, the beads were re-suspended in 1x wash buffer, and transferred to a 96-well microplate for readout in a BIORAD ZE5, and the concentration of each analyte was calculated with the standards using the LEGENDPlex data analysis software.

### Immunoblotting

Immunoblots were performed using a standard method based on our previous studies (*74*). In brief, protein samples were diluted with 2x or 4x Laemmli protein sample buffer (Bio-Rad, Cat# #1610747) followed by 100 °C boiling for 10 min, and loaded to 15 well 4%-15% Mini-PROTEAN® TGX^TM^ Precast protein gels (Bio-Rad) for standard sodium dodecyl sulfate-polyacrylamide gel electrophoresis (SDS-PAGE), Western blotting, and an enhanced chemiluminescent (ECL) substrate (ThermoFisher, Cat# 32106) or an ultra-sensitive Lumigen ECL Ultra substrate for low-femtogram-level detection (LUMIGEN, Cat# TMA-100) were applied. Primary antibodies used were diluted in 5% BSA. An anti-rabbit IgG, HRP-linked secondary antibody (Cell Signaling Technology, Cat#7074) and anti-mouse IgG, HRP-linked secondary antibody (Cat#7076) were used at a dilution of 1:5000 or 1:10000 in 5% fat-free milk. Protein band detection was performed using a ChemiDoc MP Imaging System (Bio-Rad). The band density was quantified by Image J.

### Immunoprecipitation (IP) and Streptavidin-Biotin pulldown

HEK293T/HeLa cells were transfected with expression plasmids using TransIT-X2 Dynamic Delivery System for 24 h. In some cases, cells were then treated with IFN-γ and/or transfected with LPS. The IP procedure was performed based on manuals of different magnetic beads. Briefly, whole cell lysates (WCL) were prepared in an IP lysis buffer (150□mM NaCl, 50□mM Tris, pH 7.5, 1□mM EDTA, 0.5% NP40, 10% glycerol, protease inhibitors) and incubated with 30□µL of Anti-FLAG® M2 (Millipore, Cat # M8823) or Anti-c-Myc magnetic beads (50% v/v) (ThermoFisher, Cat # 88842) rotating overnight at 4□°C. The beads were washed 3 times in 1xTBS or 5xTBST buffer according to user manuals and bound proteins were eluted with a 3x FLAG or Myc peptide.

For Streptavidin-Biotin pulldown, HeLa cells in 10-cm dishes were primed with 10 ng/ml IFN-γ for 12 h and then transfected with Biotin or Biotin-labeled LPS for 5 h. Cells were collected by centrifugation after trypsinization and lysed in nonionic detergent buffer (50 mM Tris-HCl pH7.4, 150 mM NaCl, 5 mM MgCl2, 0.1% Triton X-100) on ice for 15 min. The lysates were centrifuged for 10 min at 4□°C, 6000 g. The supernatants were served as WCL and incubated with Streptavidin Magnetic Beads (New England Biolabs, Cat # S1420S) at room temperature with rotating for 2 h. The beads were washed 3 times in binding/wash buffer (20 mM Tris-HCl, pH7.5, 0.5 M NaCl, 1 mM EDTA in ultrapure water), and boiled in SDS-PAGE reducing sample buffer for 10 min to elute biotinylated LPS with target proteins. The IP samples were subjected to immunoblotting.

### IP-MS/MS analysis

Caspase-4 and UBXN1 were overexpressed in IFN-γ-primed human cells by transfection of FLAG-CASP4 with or without Myc-UBXN1 plasmids for 24□h. The vector transfection was included as negative control. Cells were then washed by ice-cold PBS and lysed using IP lysis buffer (150□mM NaCl, 50□mM Tris, pH 7.5, 1□mM EDTA, 0.5% NP40, 10% glycerol, 1X proteinase inhibitor in di-water) on ice for 30□min with gentle shaking. The lysates were centrifuged at 14000□g, 4□°C for 15□min. The supernatants were collected as whole cell lysates and subjected to immunoprecipitation (IP) using anti-Flag M2 magnetic beads. The pulldown quality control was performed by running a Coomassie blue gel. 20□µl of the whole cell lysates were supplemented with Laemmli sample buffer, boiled for 5□min at 100□°C and stored at –20□°C as the input control. On beads samples from IP were then subjected to Ultra performance liquid chromatography-tandem mass spectrometer (UPLC-MS/MS) at the UConn Proteomics & Metabolomics Facility for acquiring and analyzing source MS/MS spectra data (Database: full Uniprot human reference proteome plus a common contaminants database from MaxQuant). Bioinformatic analyses were performed for enrichment analysis, caspase-4 bound proteins with mean >3 fold change of Average Precursor Intensity in FLAG-CASP4 group over vector control were used in analysis of binding Z-scores of caspase-4 bound proteins. The covalent ubiquitination of caspase-4 was analyzed using the method of normalizing diGly peptides to total peptides of modified and non-modified lysine residues in caspase-4.

### Immunofluorescence microscopy

Immunofluorescence staining was performed using our established methods. Cells cultured and treated in 8-well chambered microscopy slides (Ibidi, Gräfelfing, Germany, Cat # 80826) as experiment requested were sequentially wash with ice-cold PBS twice, fixed with 4% paraformaldehyde for 15 min at RT. Cells were then washed twice with cold PBS for 5 min per wash and permeabilized with 0.25% Triton X-100 in PBS (PBST) for 10 min at RT. Permeabilization solution was removed and cells were washed twice and blocked with a blocking buffer (10% goat serum, 0.1% bovine serum albumin, 0.3M glycine in PBST) for 1 h at RT. Primary antibodies (anti-CASP4/11, 1:200, R&D Cat# NB120-10454; anti-UBXN1, 1:200, Proteintech Cat# 16135-1-AP) were diluted in antibody buffer (1% BSA, 0.3M glycine in PBST) and incubated with cells overnight at 4□°C without light. Cells were washed 3 times in cold PBS for 5 min per wash, and then incubated with A mix containing a variety of Alexa Flour™ - conjugated secondary antibodies (ThermoFisher, 1:200 dilution in antibody buffer) for 1□h in the dark at RT. After 3-time wash, the nuclei were counterstained with DAPI (4’,6-diamidino-2-phenylindole, ThermoFisher, Cat # D1306, 1:1000 in antibody buffer) for 10 min in dark at RT. Cells were then washed once in PBS and imaged in PBS with a Zeiss 880 laser scanning confocal microscope (objective x 63 oil lens). All microscopic images were processed in the Zen 2.3 microscopy software (Zeiss Group, Germany) before histogram analysis in ImageJ.

### Unanchored polyubiquitin chain assay

Human cells cultured in 10-cm dishes were overexpressed with FLAG-Caspase-4 or 11 with Myc-UBXN1 by transfection of FLAG-CASP4/11 and Myc-UBXN1 plasmids for 24 h. Vector and Myc-UBXN1 transfection alone were served as controls (**Fig. 7e**). Cells were lysed with gentle shaking for 30 min in ice-cold IP lysis buffer containing 150□mM NaCl, 50□mM Tris, pH 7.5, 1□mM EDTA, 0.1% triton X-100, 10% glycerol, 1X proteinase inhibitor in di-water. The lysates were then centrifuged at 14000 g, 4□°C for 15 min, and the supernatants in each group were collected and equally split into two vials for IP with anti-Myc beads and anti-FLAG beads simultaneously. The cell lysates and IgG-conjugated beads mix were incubated with rotating overnight 4□°C. The beads were washed 3 times and split equally into two parts, one for regular elution and the other for further incubation with 0.2% SDS or USP5 (isopeptidase T, IsoT) to disrupt regular protein-protein interaction or digest unanchored free polyubiquitin chains. The USP5 eluate were collected, and beads were washed and eluted with 3X peptide to obtain caspase-4/11 bound proteins. The samples were then diluted using 4x Laemmli buffer and boiled for 10 min at 100 °C, examined by immunoblots with specific antibodies.

### Caspase-4/11 activation assay using an *in vitro* cell-lysate system

HeLa cells in 10-cm dishes were primed with 20 ng/ml of IFN-γ overnight and washed twice in cold PBS. Cells in two dishes were collected into one 1.5 ml EP tube using 500 µl of detergent free buffer (150 mM NaCl, 50 mM Tris-HCl, 1 mM EDTA, 10% glycerol, 1X protease inhibitors) by a disposable cell lifter. The cells in detergent free buffer were lysed by douncing 45∼50 times on ice using a syringe with 25G needle. The lysates were centrifuged for 15 min at 14000 g, 4□°C to remove the debris. For per reaction in general, 10 µl of the supernatants were mixed with 10 µl of various concentration of LPS and LPS mutant variants or PBS as negative control. For USP5 and USP5 inhibitor, 10 µl of the supernatants were incubated with 2 µl of USP5 (50 ng/µl) for 1 h at 30□°C, and then 10 µl of Re-LPS and 3 µl of USP5 (50 ng/µl) were added into the reaction for 4 h at 37□°C. The samples were subsequently mixed with 8 µl of 4x Laemmli protein sample buffer (Bio-Rad, Cat# #1610747) and boiled for 10 min for immunoblotting.

### *In vitro* caspase-4 activity assay

Caspase-4 activity assays in vitro with recombinant human proteins were performed in a 384-well black plate. The potential endotoxin contamination of recombinant UBXN1 proteins purified from E. coli were eliminated by using a Pierce™ High-Capacity Endotoxin Removal Spin Columns kit (ThermoFisher, Cat# 88273). In brief, recombinant caspase-4 (1 µM), UBXN1 (1 µM), K48 or K63 Tetra Ub (1 µM) and Ra-LPS (0.5 µM) were mixed in HEPES buffer (25 mM HEPES, 150 mM NaCl, 2 mM DTT, 0.5 mM EDTA pH7.5 in di-water). After brief shaking and centrifuging, 30 µl of the mixture were added to 30 µl of the Ac-LEVD-AMC caspase-4 substrate (fluorogenic, 5 mM in DMSO, Enzo, Cat# ALX-260-083-M001) in each well of the 384-well plate, protecting from light. The reaction kinetics were recorded continuously every 4 min by a Microplate Reader. The concentration of each component used in this reaction system was determined by a preliminary experiment including various concentrations.

### *In vitro* binding assay for recombinant UBXN1 protein

FLAG-Caspase-4 protein was overexpressed in HEK293T cells in 4 of 10 cm dishes by transfection of FLAG-CASP4 plasmids for 24 h. 40 µl of anti-FLAG M2 magnetic beads per dish were used to bind FLAG-caspase-4 protein in the cell lysates. The FLAG-caspase-4-beads complex were considered as on-beads FLAG-caspase-4. The on-beads samples were then split into 5 of 1.5 cm EP tubes, equal volume of anti-FLAG M2 beads incubation with lysates from vector cells were served as control. The FLAG-caspase-4-beads complex were incubated with recombinant human UBXN1 (4 µM) and commercial K48 Tetra Ub (1µM) or K63 Tetra Ub (1µM) (see “Reagent and antibodies”) in HEPES buffer (25 mM HEPES, 150 mM NaCl, 2 mM DTT, 0.5 mM EDTA pH7.5 in di-water) at 37 °C for 4 h.

### Graphing and statistical analyses

A Graphpad Prism10 software was used for graphing and statistical analyses. The sample sizes for animal experiments were estimated according to our prior experience in similar experiments. Survival curves were analyzed using a log-rank (Mantel–Cox) test. A two-tailed, unpaired Student’s *t* test or non–parametric/parametric Mann–Whitney U test was applied to a data set statistical analysis depending on its data distribution. The one-way or two-way ANOVA test was applied to simultaneously comparing multiple groups or with multiple time points. Data were presented in mean ± S.E.M, *p* values ≤0.05 were considered significant. The sample sizes, statistical tests, and *p* values were specified in figure legends.

## Data availability

The datasets generated during and/or analyzed during the current study are in the supplemental materials. Source data are provided with this paper.

## Acknowledgements

We thank Ziming Cao at UConn Health for graphing the Binding Z-scores data. This work was supported by the National Institutes of Health of Unites States grants R01AI132526 and R21AI170989 to P.W. and a UConn Health startup fund to P.W.

## Author contributions

D.Y. performed most experimental procedures, acquired and analyzed the data. J.G.C, T.G., C.W., E.T., and A.G.H assisted D.Y. with some experimental procedures and / or provided technical support. Y.W. provided key equipment and technical support. A.T.V., V.R., and J.R. contributed to data interpretation, discussion and improved writing. D.Y. and P.W. conceived, designed the study and wrote the manuscript. All the authors reviewed and/or modified the manuscript.

