## Supplemental figures and tables for "Unanchored ubiquitin chains promote the non-canonical inflammasome via UBXN1"

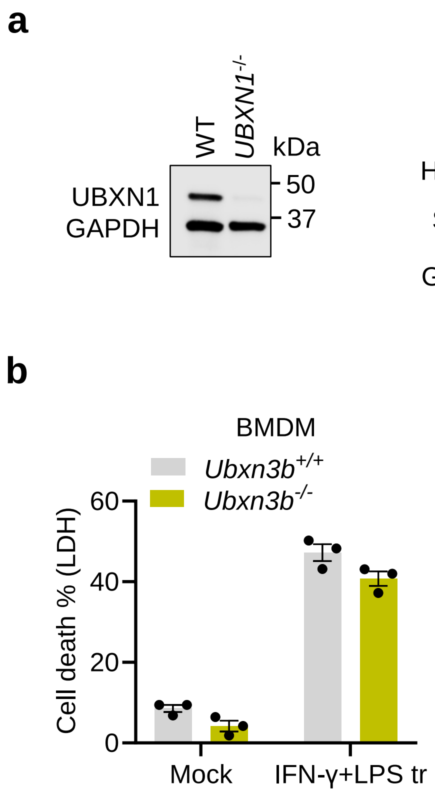

**Supplemental Fig.S1**. **Differential roles of caspase-4 binders in the non-canonical inflammasome activation. a)** Validation of *UBXN1, HUWE1*, *STUB1* and *CASP4* knockout HeLa cell by immunoblotting. **b)** Cell death of *Ubxn3b*$^{+/+}$ and *Ubxn3b*$^{-/-}$ bone marrow-derived macrophages (BMDMs) primed with 10 ng/mL of mouse IFN-γ for 3 h followed by LPS (1 µg/mL) transfection for 16 h.

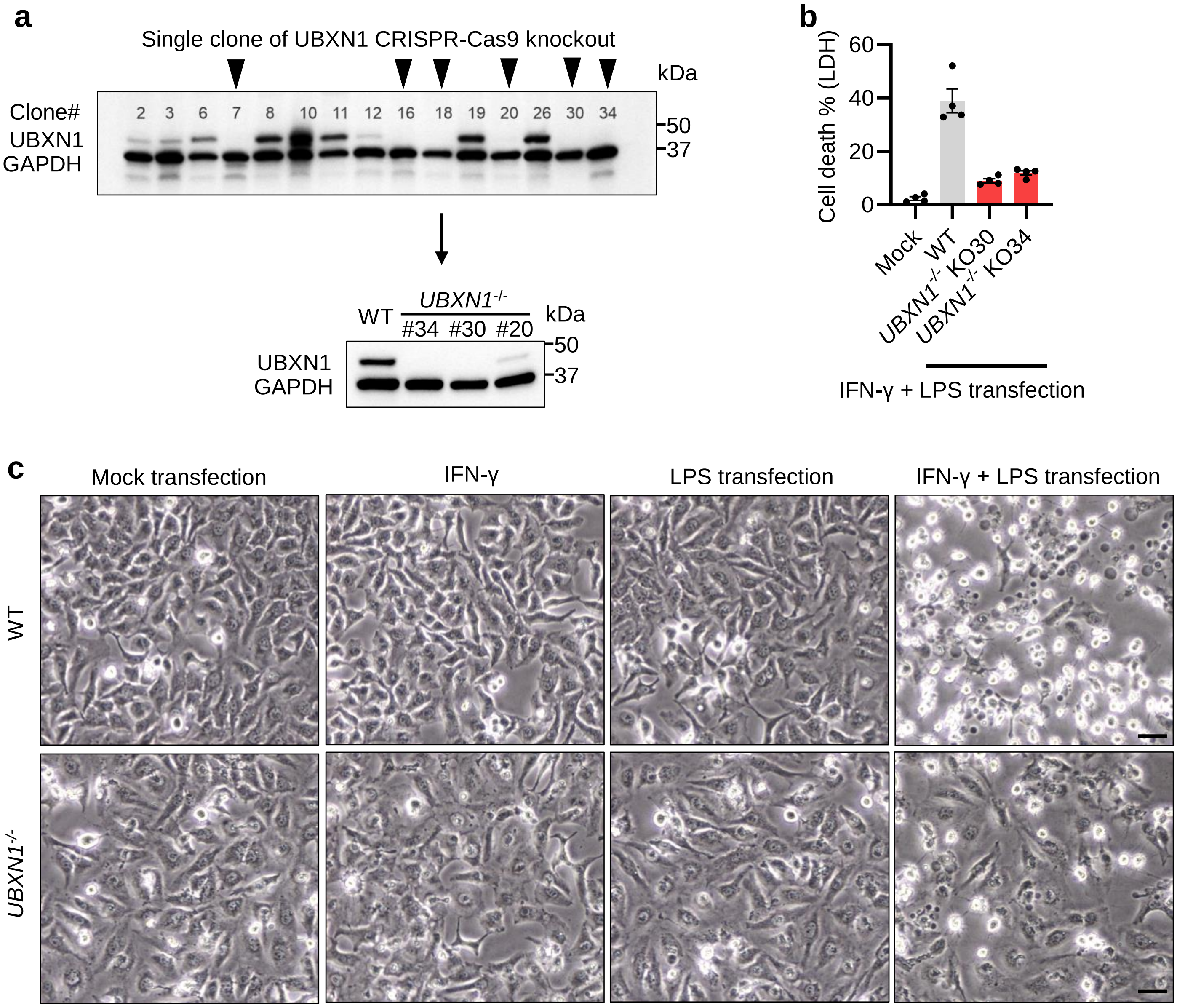

**Supplemental Fig.S2**. **UBXN1 is critical for non-canonical inflammasome signaling in human cells.** **a)** Validation of monoclonal UBXN1 knockout HeLa cell by immunoblotting. **b)** Percent cell death induced by human IFN-γ (10 ng/mL) overnight plus 1 μg/mL of LPS transfection for 5 h. **c)** Differential interference contrast (DIC) microscopy of HeLa cells treated as in **(b)**. Bright, round cells are typical dying cells. Scale bar: 20 μM.

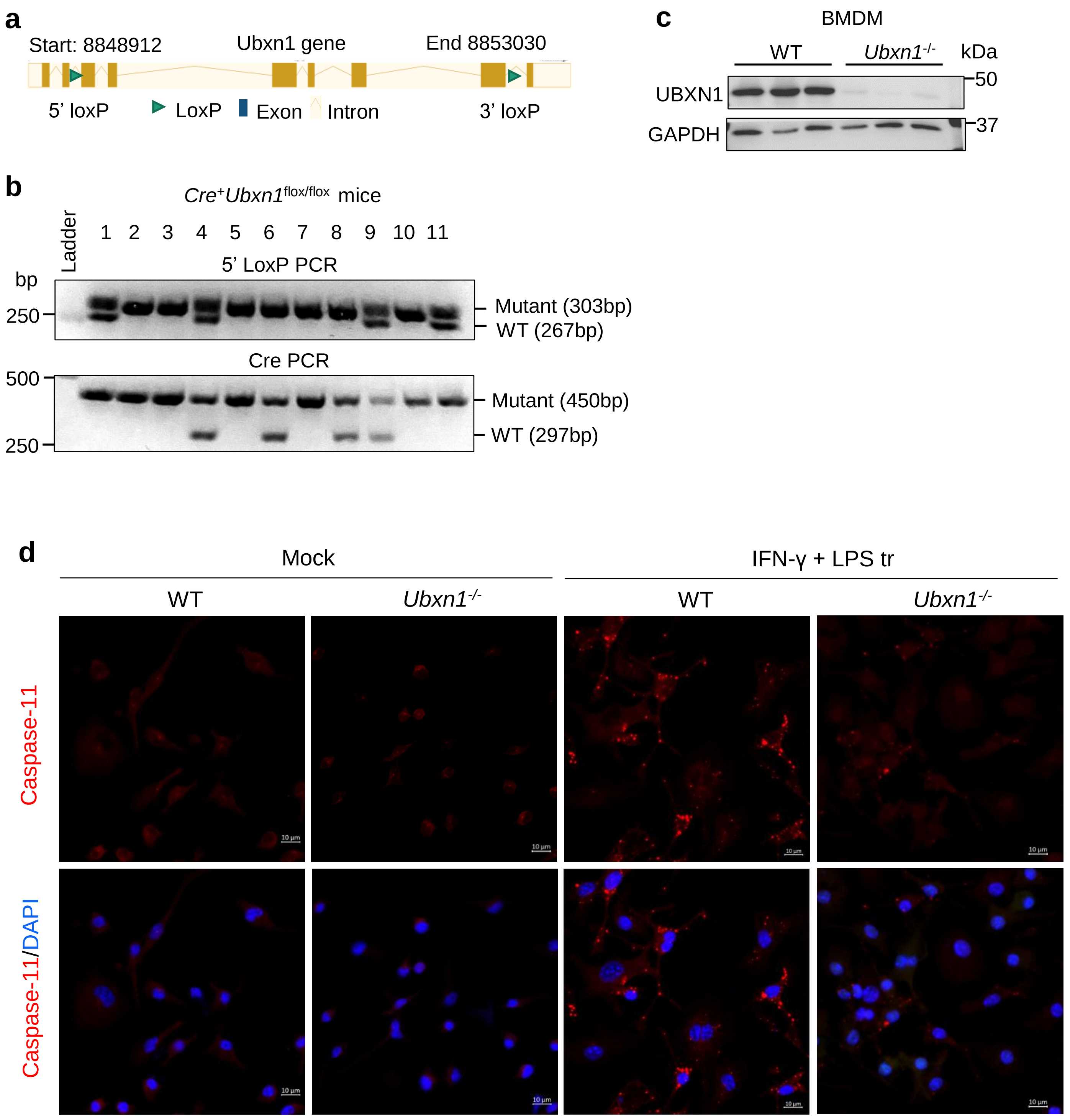

**Supplemental Fig.S3**. **UBXN1 is crucial for non-canonical inflammasome activation in mouse macrophages. a)** A graph depicting *Ubxn1* gene structure and insertion of loxP. **b)** Genotyping PCR of Cre$^{+}$*Ubxn1*$^{flox/flox}$ mice. **c)** Immunoblots of UBXN1 protein expression in bone marrow-derived macrophages (BMDMs). **d)** Immunofluorescence staining of caspase-11 in WT and *Ubxn1*$^{-/-}$ BMDMs treated with 10 ng/mL of mouse IFN-γ for 3 h plus LPS (transfection,1μg/mL) for 16 h. LPS tr: LPS transfection, Mock: transfection reagent only.

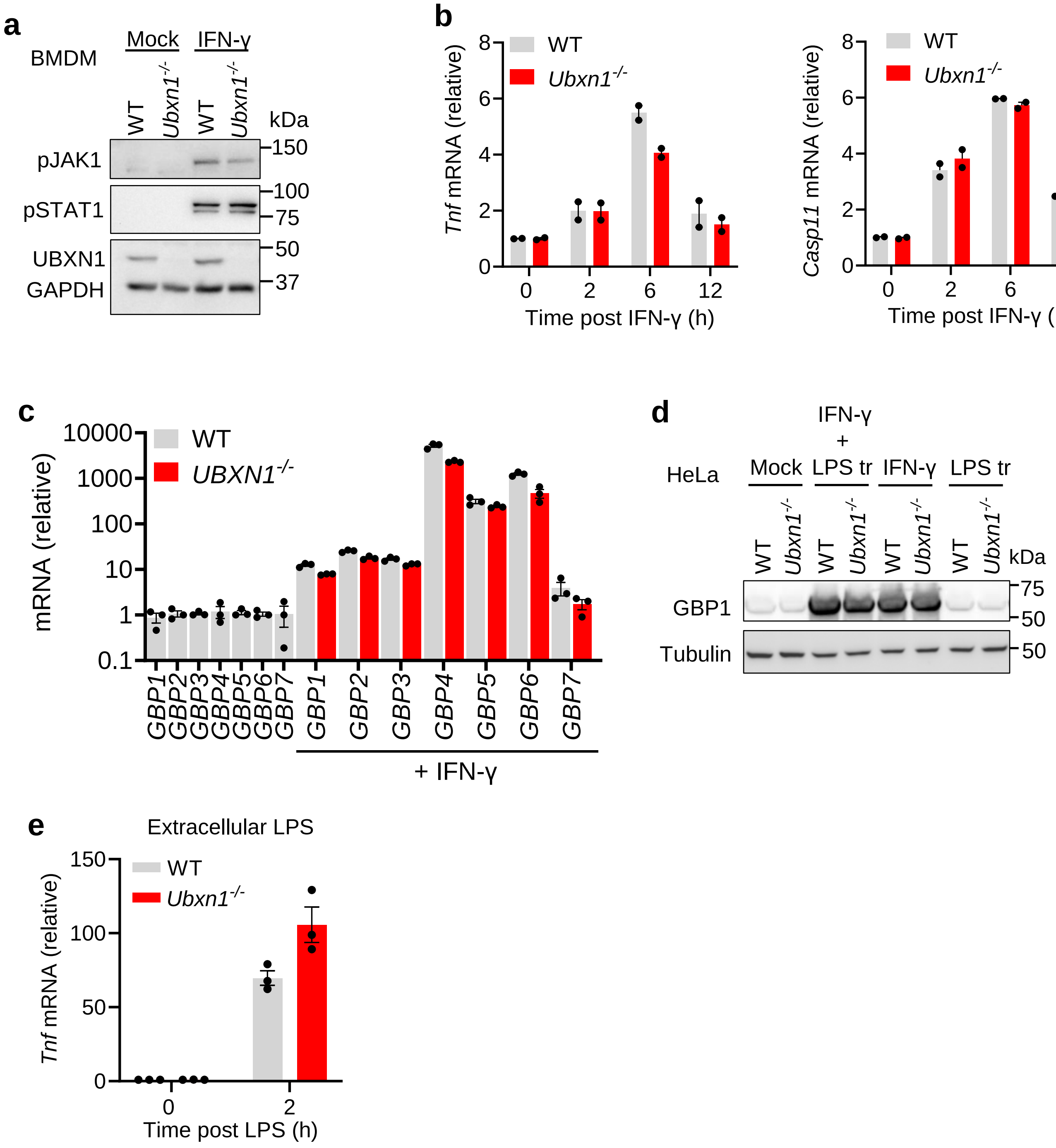

**Supplemental Fig.S4. UBXN1 is dispensable for IFN-γ-JAK-STAT and TLR4 signaling**. **a)** Phosphorylation of JAK1 and STAT1 in primary bone marrow-derived macrophages (BMDMs) treated with 10 ng/mL of mouse IFN-γ for 30 min. **b)** Quantitative RT-PCR of mRNA expression for the indicated genes in BMDMs treated with 10 ng/mL of mouse IFN-γ. **c)** The expression of GBPs in WT and *UBXN1*$^{-/-}$ HeLa cells treated with human IFN-γ for 12 h. **d)** GBP1 expression in HeLa cells primed with 10 ng/mL of IFN-γ for 12 h, and then transfecetd with 1 µg/mL of LPS for 5 h. **e)** The *Tnf* mRNA expression in BMDMs stimulated with 0.5 µg/mL of LPS. Bar: mean ± SEM. Each symbol represents one mouse. Mock: transfection reagent only.

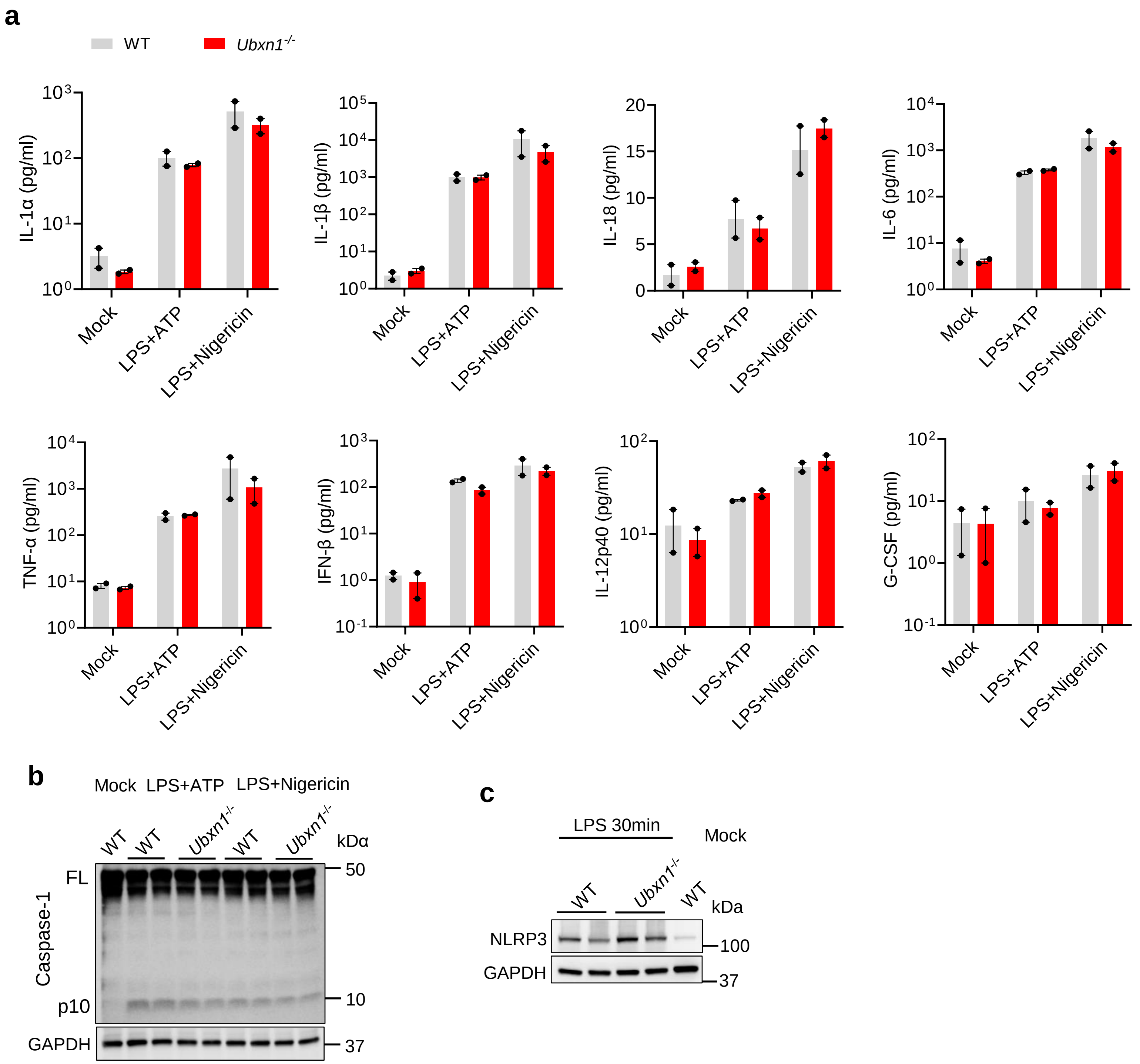

**Supplemental Fig.S5. UBXN1 is dispensable for the canonical NLRP3 inflammasome signaling**. **a)** Primary bone marrow-derived macrophages (BMDMs) were primed with LPS (1 μg/ml) for 3 h and then stimulated by ATP (5 mM) or Nigericin (10 μg/ml) for 30 min. The concentrations of cytokines/growth factors in the cell culture medium were analyzed by a bead-based multiplex ELISA. Bar: mean ± SEM. N=2 mice/group. **b)** The immunoblots of caspase-1 in BMDMs treated as in **a)**. Each lane = one mouse. FL: full-length. **c)** The immunoblots of NLRP3 in BMDMs primed with 1 μg/mL of LPS for 30 min. Mock: phosphate buffered saline.

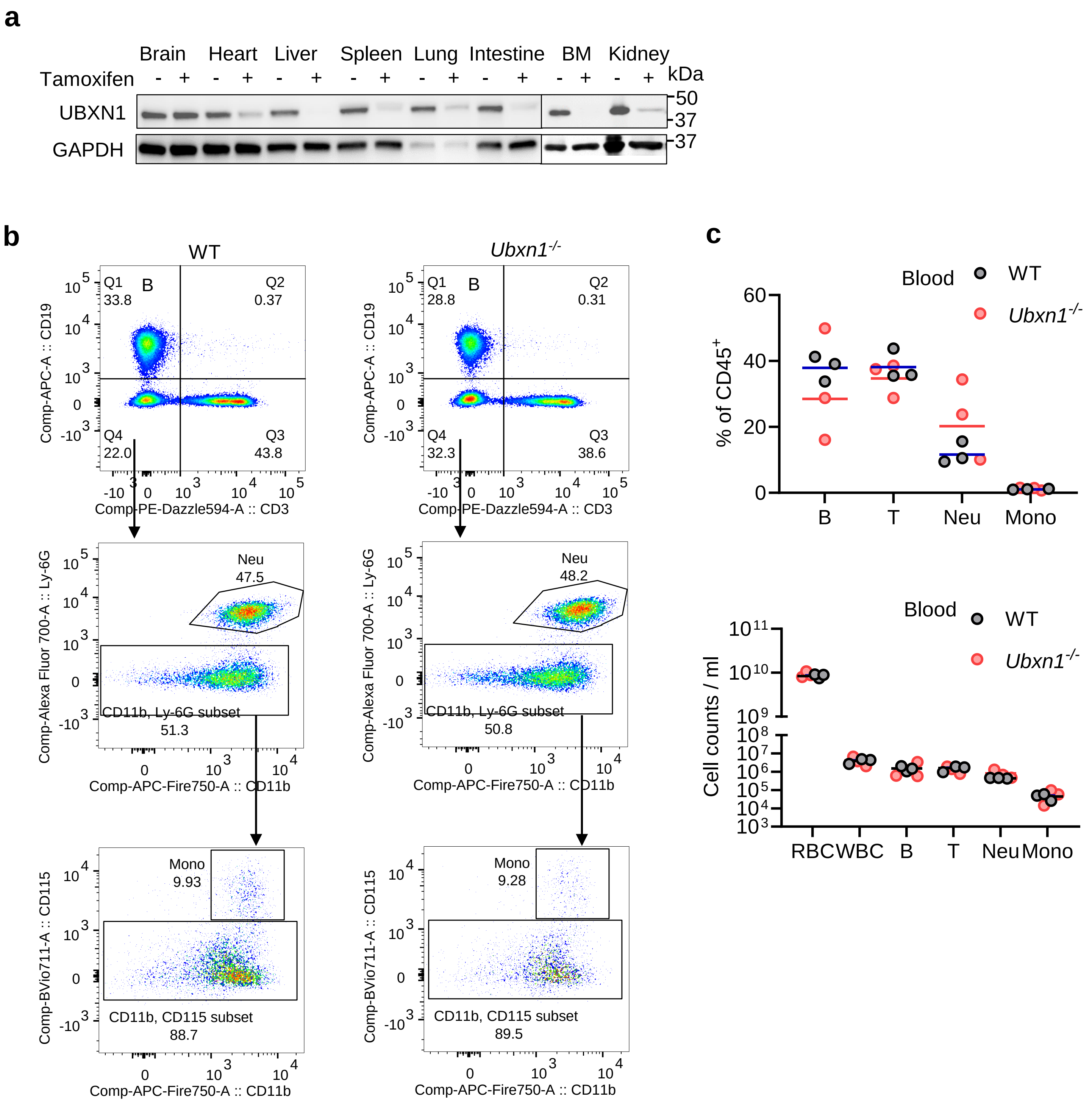

**Supplemental Fig.S6. Normal peripheral immune populations in *Ubxn1*$^{-/-}$ mice. a)** Immunoblots of UBXN1 protein in various tissues from the Cre$^{+}$*Ubxn1*$^{flox/flox}$ mice treated without (-) or with tamoxifen (+). BM: bone marrow. **b)** The gating strategy and **(c)** ratios and counts of major blood immune populations to total leukocytes (CD45$^{+}$). The blood immune populations in aged- and sex-matched mice were quantitated by flow cytometry with specific surface markers. RBC: red blood cell , WBC: white blood cell CD45, T cell: CD3, B cell: CD19, Neutrophil: Ly6G, Monocyte: CD115/CD11b. Bar: mean ± SEM, N=3 mice/group.

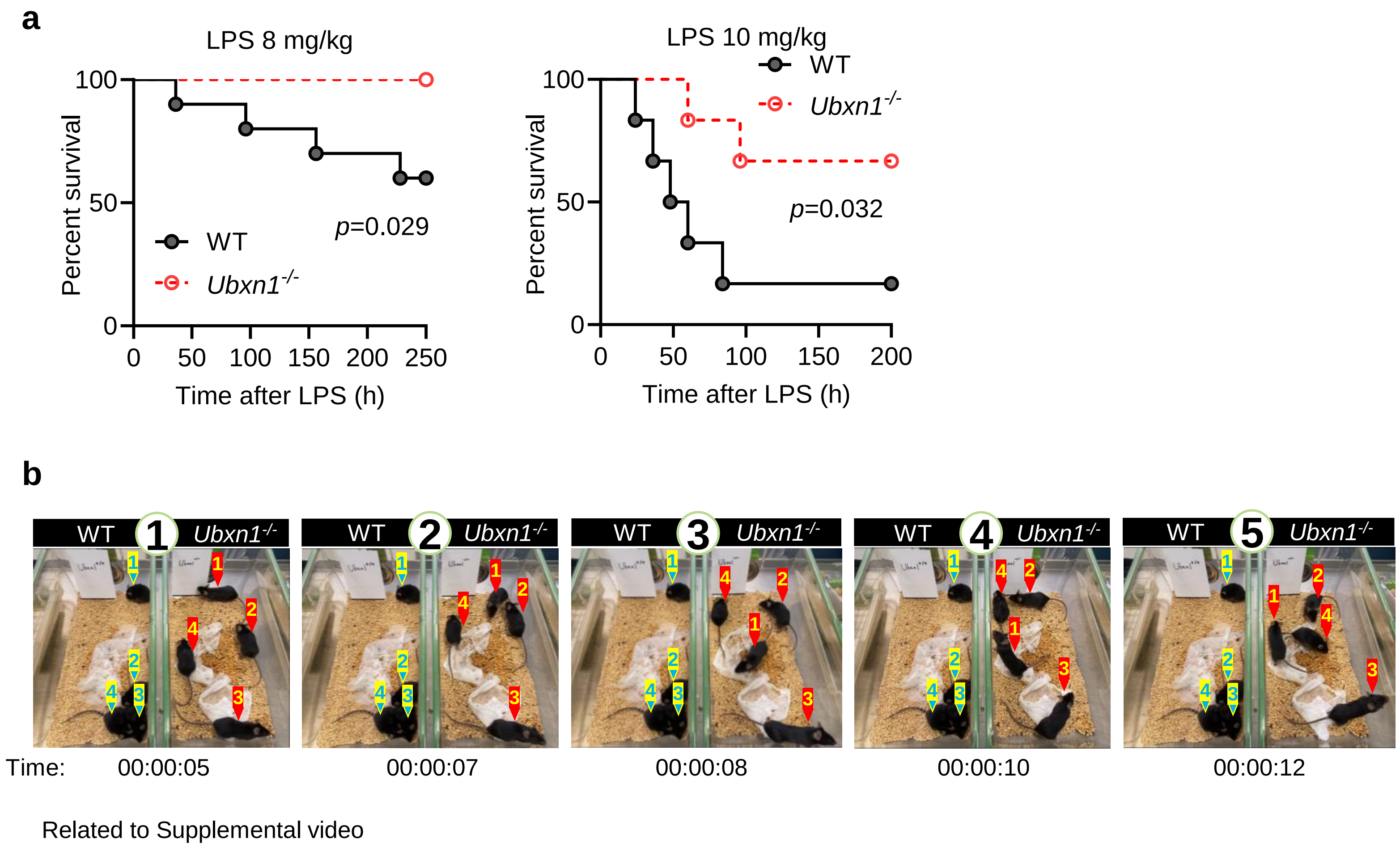

**Supplemental Fig.S7. *Ubxn1*$^{-/-}$ mice are resistant to lethal LPS sepsis. a)** The survival curves of age- and sex-matched littermates administered with the indicated dose of LPS intraperitoneally. For 8 mg/kg of LPS, N=10 for WT, and *Ubxn1*$^{-/-}$ respectively, *p*=0.029; for 10 mg/kg of LPS, N=6 for WT, and *Ubxn1*$^{-/-}$ respectively. *p*=0.032, Log-Rank test. **b**) Time-lapse morbidity of age- and sex-matched littermates injected with LPS (10 mg/kg) intraperitoneally for 36 h. Related to Supplemental video.

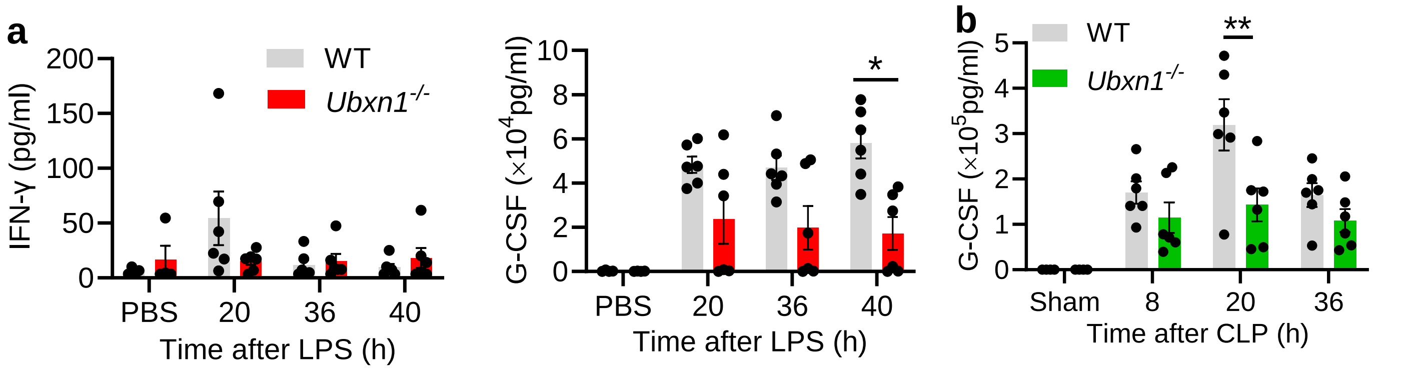

**Supplemental Fig.S8**. **UBXN1 amplifies systemic inflammatory responses during sepsis**. Age- and sex-matched littermates were injected with 10 mg/kg of LPS intraperitoneally or subjected to a cecal-ligation-and–puncture (CLP) procedure. The concentrations of cytokines/growth factors in the **a**) sera (at 20, 36 and 40 h after LPS), and **b**) sera at 8, 20, 36 h post CLP were analyzed by a bead-based multiplex ELISA. Bar: mean ± s.e.m. Each dot = one animal. *$p$<0.05, **$p$<0.01 (Two way-ANOVA, Dunnett comparisons).

**a**

Kn-Ub mutants

| | 6 | 11 | 27 | 29 | 33 | 48 | 63 |
|---|---|---|---|---|---|---|---|
| HA-Ub | K | K | K | K | K | K | K |
| HA-K6 | K | R | R | R | R | R | R |
| HA-K11 | R | K | R | R | R | R | R |
| HA-K27 | R | R | K | R | R | R | R |
| HA-K29 | R | R | R | K | R | R | R |
| HA-K33 | R | R | R | R | K | R | R |
| HA-K48 | R | R | R | R | R | K | R |
| HA-K63 | R | R | R | R | R | R | K |

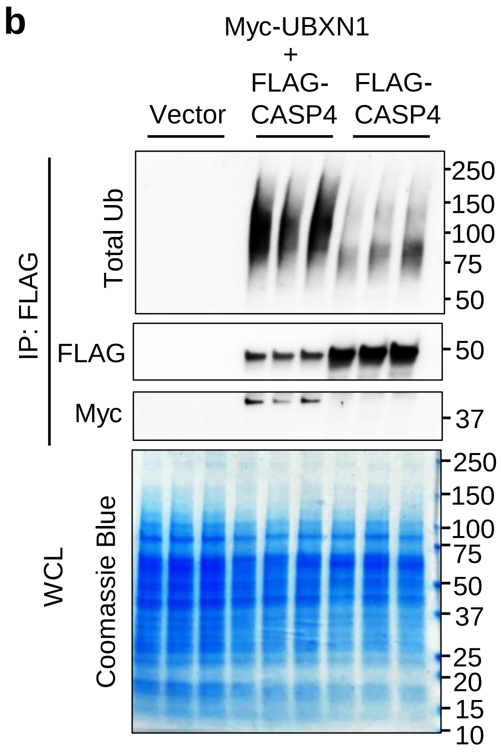

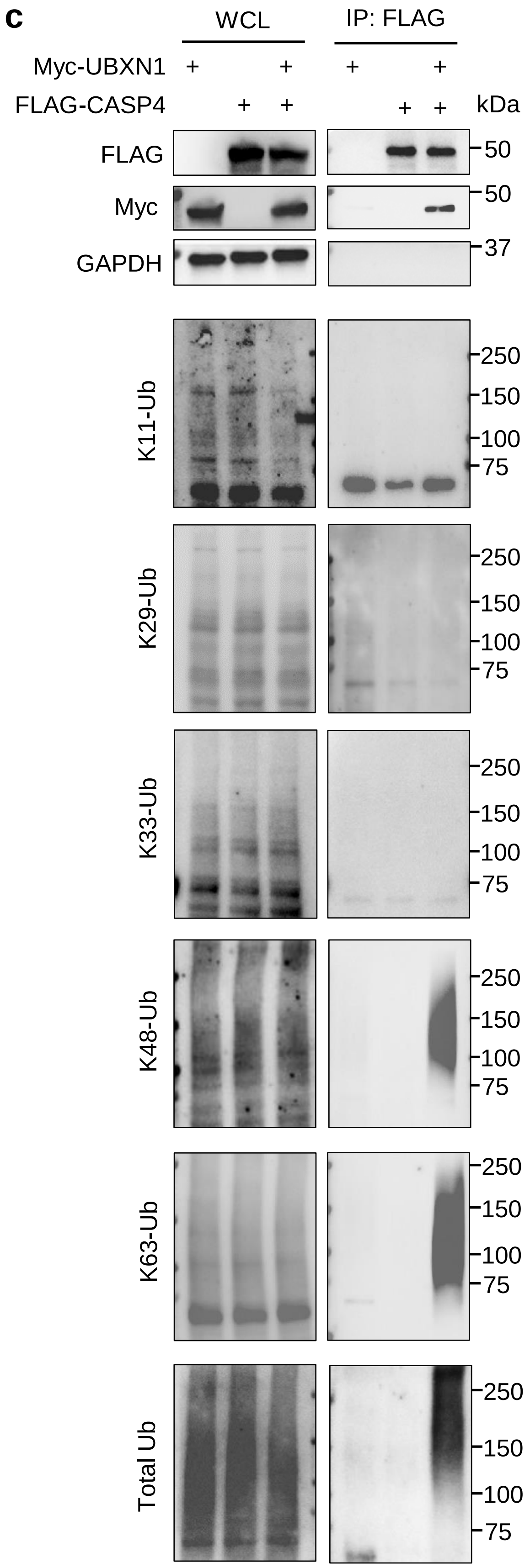

**Supplemental Fig.S9**. **UBXN1 enhances K48- and K63- linked ubiquitination of caspase-4**. **a)** A graph illustrating mutant Kn-Ub. **b, c)** Co-immunoprecipitation (IP) of FLAG-CASP4 and bound Myc-UBXN1 with an anti-FLAG antibody from HEK293T cells transfected with different combinations of FLAG-CASP4, Myc-UBXN1 and vector plasmids. Shown are immunoblots and Coomassie blue staining. Endogenous Ub is detected with a polyubiquitin (total Ub) or linkage specific Ub antibody as indicated. WCL: whole cell lysate.

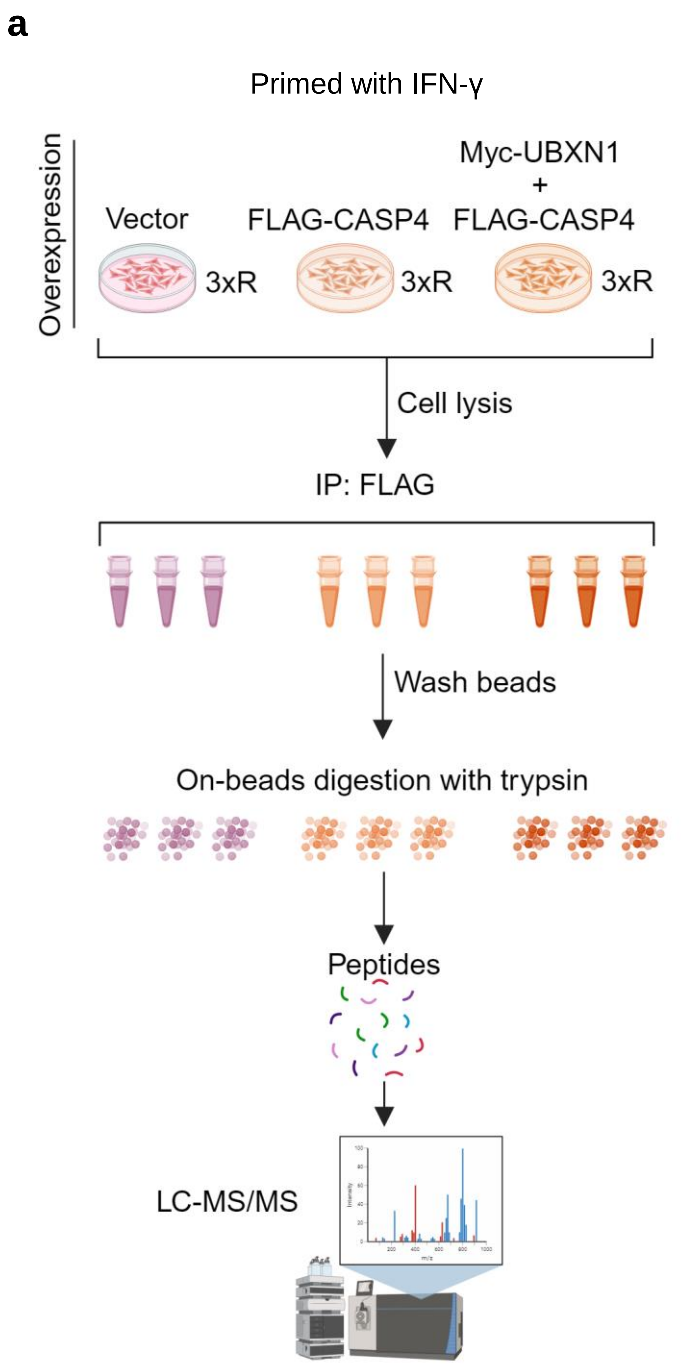

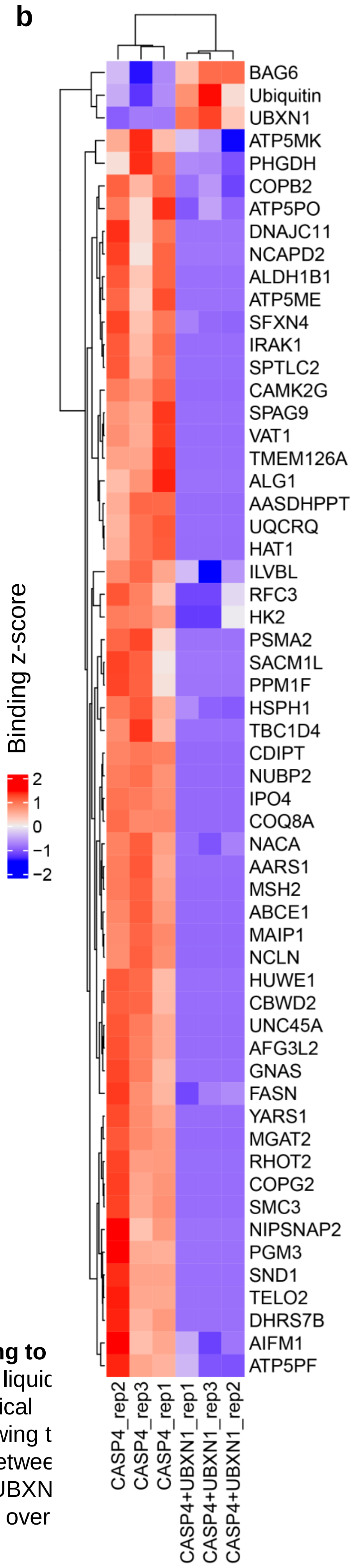

**Supplemental Fig.S10**. **UBXN1 promotes ubiquitin binding to caspase-4. a)** The workflow of immunoprecipitation (IP) and liquic chromatography- mass spectrometry (LC-MS). 3xR: 3 biological replicates. Created with BioRender.com. **b)** A heat map showing t binding Z-scores of 58 proteins with significant differences betwee FLAG-caspase-4 + Myc vector and FLAG-caspase-4+ Myc-UBXN groups. $p<0.05$ two-tailed Student's t-test, Log2 fold-change over vector >1.5.

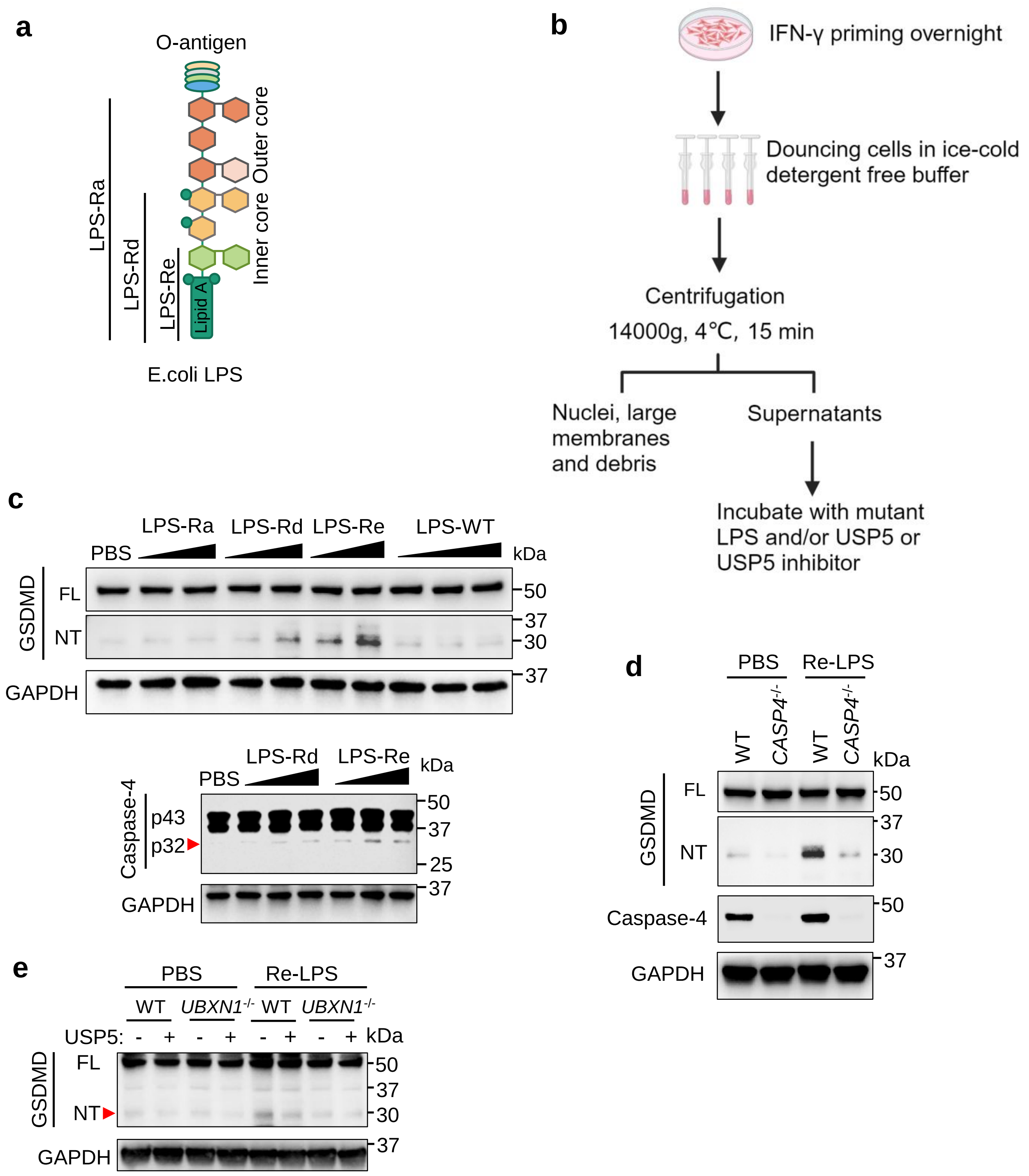

**Supplemental Fig.S11**. **Short LPS mutants activate the noncanonical inflammasome in cell-free lysates.** **a)** A schematic illustration of *E.coli* WT, Ra, Rd and Re mutant LPS structures. **b)** The procedures to obtain cell free lysates for studying the noncanonical inflammasome. Created with BioRender.com. **c)** The immunoblot of GSDMD and caspase-4 in IFN-γ-primed HeLa cell lysates treated with increasing doses of LPS or mutants for 4 h at 37 ℃. **d)** The immunoblots of GSDMD and caspase-4 in IFN-γ-primed WT and *CASP4*$^{-/-}$ cell lysates upon Re-LPS stimulation. **e)** The immunoblots of GSDMD in IFN-γ-primed WT *and UBXN1*$^{-/-}$ HeLa cell lysates incubated with USP5 and Re-LPS or PBS (phosphate buffered saline). NT; N-terminal fragment, FL: full-length.

| Supplemental Table 1 | |
|---|---|
| **Sequences for gRNAs for CRISPR-Cas9 knockout of genes in human cell lines** | |
| *UBXN1*-pair1 | Forward: CACCGTCTAAAGGCTCGTCCACATC |
| | Reverse: AAACGATGTGGACGAGCCTTTAGAC |
| *UBXN1*-pair2 | Forward: CACCGACACTGGTCATACTCCCGCT |
| | Reverse: AAACAGCGGGAGTATGACCAGTGTC |
| *STUB1* | Forward: CACCGGGCCGTGTATTACACCAACC |
| | Reverse: AAACGGTTGGTGTAATACACGGCC |
| *HUWE1* | Forward: CACCGTTCCATCGAAGCGGTCCAAC |
| | Reverse: AAACGTTGGACCGCTTCGATGGAA |
| *CASP4* | Forward: CACCGGAGAAACAACCGCACACGCC |
| | Reverse: AAACGGCGTGTGCGGTTGTTTCTCC |

| **Supplemental Table 2** | |
|---|---|
| **Primers for RT-qPCR** | |
| mouse *Tnf* | forward: GGTGCCTATGTCTCAGCCTCTT |
| | reverse: GCCATAGAACTGATGAGAGGGAG |
| mouse *Casp11* | forward: GTGGTGAAAGAGGAGCTTACAGC |
| | reverse: GCACCAGGAATGTGCTGTCTGA |
| Human *GBP1* | forward: TCAATGAGGAAATCCCAGCCC |
| | reverse: AGGCTGTTCCCTTGTCTGTTC |
| Human *GBP2* | forward: ATCTCTGATCTGGGGAACAACAC |
| | reverse: GATAGAGGCCCACAATCGCC |
| Human *GBP3* | forward: AGCACAGACAAGAGAACAATGCC |
| | reverse: TCTGGATTCGCCACCAGTTC |
| Human *GBP4* | forward: CAGTGCCCACACCAGGTTATC |
| | reverse: TTCCTGTGCGGTATAGCCCT |
| Human *GBP5* | forward: CGGCGATTCAAAGGCAGAAC |
| | reverse: AGCCTGTTCCTGCATCTGTTG |
| Human *GBP6* | forward: ACTGCACCATCCCATTTGTGG |
| | reverse: TGCCAACCTAGAAGAGCCTGC |
| Human *GBP7* | forward: ACTCTGGACAGAGGAACGCC |
| | reverse: TAGAGGCCCACAATTGCCAC |
